# Most autophagic cell death studies lack evidence of causality

**DOI:** 10.1101/2024.12.22.629975

**Authors:** Ali Burak Özkaya, Yasmin Ghaseminejad

## Abstract

Autophagy plays a critical role in maintaining cellular homeostasis and is implicated in various physiological and pathological processes, including cancer, neurodegeneration, and metabolic disorders. Although typically associated with cell survival, autophagy has also been proposed to contribute to cell death, referred to as autophagic cell death (ACD). However, the identification of ACD remains contentious due to inconsistencies in experimental methodologies and terminological misuse.

In this study, we systematically evaluated 104 research articles published in 2022 that claimed to demonstrate ACD. Articles were assessed based on established criteria, including evidence for autophagy, evidence for cell death, exclusion of apoptosis, and experimental designs demonstrating causality. Our findings reveal that only 12.5% of the articles fulfilled all ACD criteria, while 37.5% provided only correlation-level evidence. Additionally, 54.81% failed to demonstrate autophagy flux, 32.7% relied on viability loss rather than direct evidence of cell death, and 45.0% of studies utilizing autophagy inhibition failed to demonstrate actual inhibition of autophagy. Inconsistent terminology was also prevalent, with ‘autophagy-mediated cell death’ often misclassified as ACD and ACD frequently misused to describe autophagy co-occurring with cell death.

These issues highlight a lack of rigor in current practices, with correlation-level evidence, inappropriate experimental designs, and terminological misuse undermining study robustness. To address these challenges, we developed a systematic workflow providing experimental and analytical guidance for classifying evidence for different modes of autophagy. Our analysis underscores the need for greater rigor, standardized approaches, and precise terminology to advance understanding of the interplay between autophagy and cell death.

## Introduction

Autophagy is a physiological process in which cellular components such as proteins and organelles are recycled via lysosomal activity. This process is characterized by the engulfment of the target cargo by autophagosomes, which then fuse with the lysosomes to break down complex molecules and structures into simple building blocks. Autophagy serves as both a catabolic pathway triggered as a response to starvation to provide cells with nutrients, and a cytoprotective mechanism that eliminates potentially harmful substances, such as misfolded proteins, dysfunctional mitochondria and pathogens. Due to its critical role in cellular homeostasis, disruption of the autophagy machinery, resulting in either excessive or insufficient autophagy, can lead to undesirable outcomes for the cell, such as cell death or neoplastic transformation. These disruptions can have severe consequences at the organismal level, potentially leading to malignant, neurodegenerative, or metabolic disorders [1].

The role of autophagy in cell death is complex and not as straightforward as one might expect from a cytoprotective pathway. Autophagy has been proposed as an alternative programmed cell death mechanism, where cells lose the ability to maintain themselves due to excessive self-digestion, leading to their demise [2]. However, given its apparent role in cell survival, discussions about its true role in cell death have been controversial at best [3], [4]. Excessive autophagy has been observed in many cases of cell death, raising the question of whether cells are dying by autophagy (Autophagic Cell Death, ACD) or are activating autophagy as a survival strategy in an attempt to avoid death (Autophagy-Associated Cell Death) [5]. Shen and Codogno proposed following criteria to identify autophagic cell death: increased autophagy flux, prevention of death by suppression of autophagy, and exclusion of apoptosis [6]. Similarly, the Nomenclature Committee on Cell Death (NCCD) specified that autophagic cell death should only be identified if inhibition of autophagy suppresses the cell death [7].

More recently, the terms ‘autophagy-mediated cell death’ (where excessive autophagy triggers apoptosis or another type of cell death) and ‘autophagy-dependent cell death’ (where autophagy itself is the primary cause of cell death) have been introduced to distinguish between different mechanisms and clarify the role of autophagy in these processes [8], [9]. However, since excluding other modes of cell death is one of the criteria for identifying ACD, the concept of ‘autophagy-dependent cell death’ essentially becomes synonymous with ACD [10]. Furthermore, within the spectrum of ACD, there is a specific form known as autosis, which is characterized as an autophagy-dependent, non-apoptotic cell death process with distinct morphological and biochemical features [11].

Even though it has been more than a decade since autophagic cell death was first distinguished from autophagy-associated cell death, and numerous follow-up reviews and guides have been published [10], it remains unclear whether current studies claiming to demonstrate autophagic cell death meet the established criteria and show a causal relationship between autophagy and cell death. In this study, we aimed to investigate the evidence provided in studies claiming to demonstrate ‘autophagic cell death’ and analyse how they interpret their findings.

## Materials and Methods

### 1. Article retrieval and database creation

The following search string was used in Web of Science to retrieve articles published in 2022 that claimed to demonstrate autophagic cell death: ALL=(“autophagy-associated cell death”) OR ALL=(“autophagic cell death”) OR ALL=(“type II cell death”) OR ALL=(“(type II) cell death”) OR ALL=(“autosis”) OR ALL=(“autophagy-mediated cell death”) OR ALL=(“autophagy-dependent cell death”).

Information regarding each article, including its title, DOI number, the journal in which it was published, and details of the experimental designs and methods used, was exported to an online spreadsheet database (Airtable).

### 2. Investigation of the claims and exclusion criteria

We investigated each article to determine the specific claim made by the authors—whether it was autophagic cell death, type II cell death, autophagy-mediated cell death, or autophagy-associated cell death—and identified where and in what context these claims were made. We included only those articles where the claim of autophagic cell death was relevant to the main purpose and hypothesis of the study. Therefore, all of the included articles contained an intervention strategy (such as chemical treatment or RNA interference) which authors claimed to have induced autophagic cell death. Reviews and articles that merely mentioned autophagic cell death without substantial context were excluded.

### 3. Analyses of the evidence

The evidence of autophagic cell death was analysed on multiple levels by asking various questions derived from the criteria originally proposed by Shen and Codogno [6]. We sought to answer the following questions:

1. Is there sufficient evidence for autophagy?
2. Is there sufficient evidence for cell death?
3. Is apoptosis excluded?
4. Is there an experimental design to demonstrate autophagic cell death?

We evaluated all evidence presented in each study by investigating what was measured by using which method and whether the result of this measurement is a reliable indicator of the related claim (autophagy, cell death or apoptosis) according to the available guides. When multiple measurements were carried out to support a single claim, we focused on the strongest evidence. Additionally, we assessed whether the experimental design could effectively identify autophagic cell death.

Beyond these questions, we also investigated whether the data produced by the experiments were correctly interpreted.

#### 1. Is there sufficient evidence for autophagy?

We determined the answer to this question in accordance with the ‘recommended methods for monitoring autophagy’ table presented in ‘Guidelines for the Use and Interpretation of Assays for Monitoring Autophagy’ [12]. Briefly, if visualization of autophagic vacuoles, measurement of LC3-II, and/or measurement of SQSTM1/p62 was performed, we considered it sufficient evidence for demonstrating a change in autophagy. However, other markers, including transcription-level expressional changes in Beclin1 and other autophagy-related proteins, spectrophotometric measurement of total acridine orange (and other acidotropic dye) signals as well as changes in the activity of signalling pathways such as mTOR and JNK alone, were considered insufficient.

Autophagy flux (autophagosome flux, autophagic flux) is the flow of autophagic degradation from autophagosome formation to lysosomal degradation. Unlike static markers like LC3-II or p62, which can ambiguously indicate either upregulated autophagy or stalled flux, assessing autophagy flux provides a complete view of autophagic activity [13]. Acceptable evidence of autophagy flux includes comparing LC3-II and p62 levels with and without lysosomal inhibitors (e.g., Bafilomycin A1), demonstrating differential localization in dual-labelled fluorescent reporters (e.g., mRFP-GFP-LC3), and tracking cargo degradation through time-course analyses [14].

#### 2. Is there sufficient evidence for cell death?

Results obtained from methods measuring loss of membrane integrity, such as lactate dehydrogenase assays, dye exclusion tests, and specific markers were considered sufficient evidence to demonstrate cell death. Additionally, indicators of apoptosis, including Annexin-V/PI staining, caspase activity, PARP cleavage, and DNA fragmentation were considered sufficient evidence to demonstrate apoptosis. We distinguished apoptosis from non-specific cell death, as the presence of apoptosis indicates that the death may not be autophagic.

Results obtained from methods measuring viability loss, such as formazan-based or similar metabolic activity assays, measurement of total DNA or protein content, and counting the number of cells or colonies, were considered insufficient evidence for cell death. This is because viability loss may result from decreased proliferation rates rather than increased cell death [15]. However, to distinguish viability loss from no evidence, we included it in our findings as a separate entity.

We accepted Annexin-V/PI, live/dead assays, as well as other similar dual markers as non-apoptotic indicators of cell death as PI or other dead cell indicator component of such assays measures loss of membrane integrity.

#### 3. Is apoptosis excluded?

We consider apoptosis to be excluded when its absence is proven and/or when there is an experimental approach to exclude it. Absence of apoptosis should be demonstrated via an acceptable method (listed in 3.2.) in the setting where cell death occurs. An acceptable experimental design should demonstrate that cell death persists despite the suppression of apoptosis through chemical (such as caspase inhibitors) or genetic (such as caspase siRNA) inhibition. An alternative method could be utilizing cells lacking components of apoptotic machinery such as BAX/BAX KO cells. However, we have not encountered this method in our investigation.

#### 4. Is there an experimental design to demonstrate autophagic cell death?

Experiments where suppression of autophagy prevents cell death and/or further activation of autophagy increases cell death were considered as appropriate designs to identify autophagic cell death.

We considered targeting proteins and using small molecules listed on “genetic and pharmacological regulation of autophagy table” in “Guidelines for the Use and Interpretation of Assays for Monitoring Autophagy” [12] as appropriate strategies to inhibit autophagy. Genetic regulation in this context includes both transient interference methods, such as siRNAs, and gene knockout approaches targeting autophagy-related genes. Other strategies such as small molecule inhibitors of non-specific signalling pathways and genetic targets that are not essential for autophagy were considered inappropriate.

## Results

### 1. Database Construction and Article Characteristics

We investigated a total of 137 articles from 94 different journals. After excluding 7 review articles, 21 articles that did not contain relevant claims, and 4 articles for which the full texts were not accessible, 104 articles from 76 different journals (75.9%) were included in the final analysis. Among the excluded articles, one with an ‘autophagy-associated cell death’ claim (Article ID = 10; see Supplementary Data) was omitted as it correctly used the term and provided correlation-level evidence, which does not meet the criteria for an autophagic cell death claim.

We entered all relevant information regarding these articles to an online spreadsheet (Airtable) creating a database for our analysis (See supplementary document for the complete database).

98 out of 104 (94.2%) articles contained ‘autophagic cell death’ (ACD) claim, 5 (4.8%) contained ‘autophagy-mediated cell death’, 3 (2.9%) contained ‘autophagy-associated cell death’, 3 (2.9%) contained ‘autosis’ and 2 (1.9%) contained ‘autophagy-dependent cell death’ claims. There were 7 (6.7%) articles with multiple claims in their manuscripts. It is important to note that, except for the omitted article (ID=10), in all 3 instances where authors used ‘autophagy-associated cell death’ they also used ‘autophagic cell death’ as if they were synonyms (see Supplementary data).

### 2. Experimental Models

The most commonly used experimental models were commercial cell lines, employed in 67 studies (64.42%), while the remaining studies utilized tissue samples or primary cells from various sources. Across the dataset, a total of 164 distinct commercial cell lines were used. The most frequently used were A549 and HCT116, each appearing in 9 studies, followed by MCF-7, MDA-MB-231, and HEK-293T, each used in 8 studies (Supplementary data).

### 3. Evidence for Autophagy

Surprisingly, 8 out of 104 articles (7.7%) failed to provide appropriate evidence for autophagy. Among the remaining 96 articles, which provided evidence, the vast majority (n=88) employed LC3-II measurement in combination with other methods to monitor autophagy. Other commonly used strategies included demonstrating a reduction in p62/SQSTM1 levels (n=48) and visualizing autophagic vacuoles (n=29) through various techniques (see Supplementary data).

47 articles (45.2%) provided evidence for a positive autophagy flux. The main evidence was provided by LC3-II turnover itself (n=12) or in combination with other methods (n=24). p62 degradation and rescue was also common (n=24) but only when it is combined to LC3-II turnover (n=23). Other methods include demonstrating an active autophagosome by reporter assays and visualization (n=11) but these are mainly employed as the only strategy (n=10) (See supplementary data). The details of the inhibitors used to demonstrate autophagy flux are be mentioned in the next section.

### 4. Experimental Design to Demonstrate Autophagic Cell Death

Of the 104 articles, 60 (57.7%) lacked specific experimental designs to distinguish autophagic cell death (Figure 1A). Among these, 8 provided no evidence at all, and 12 demonstrated autophagy induction alone as evidence for ACD without any supporting data. The remaining articles without specific experimental designs produced correlation-level evidence. In these studies, while demonstrating autophagy correlating to cell death, 13 articles relied on viability loss to indicate cell death, and 27 actually measured cell death (Figure 1A).

**Figure 1.**
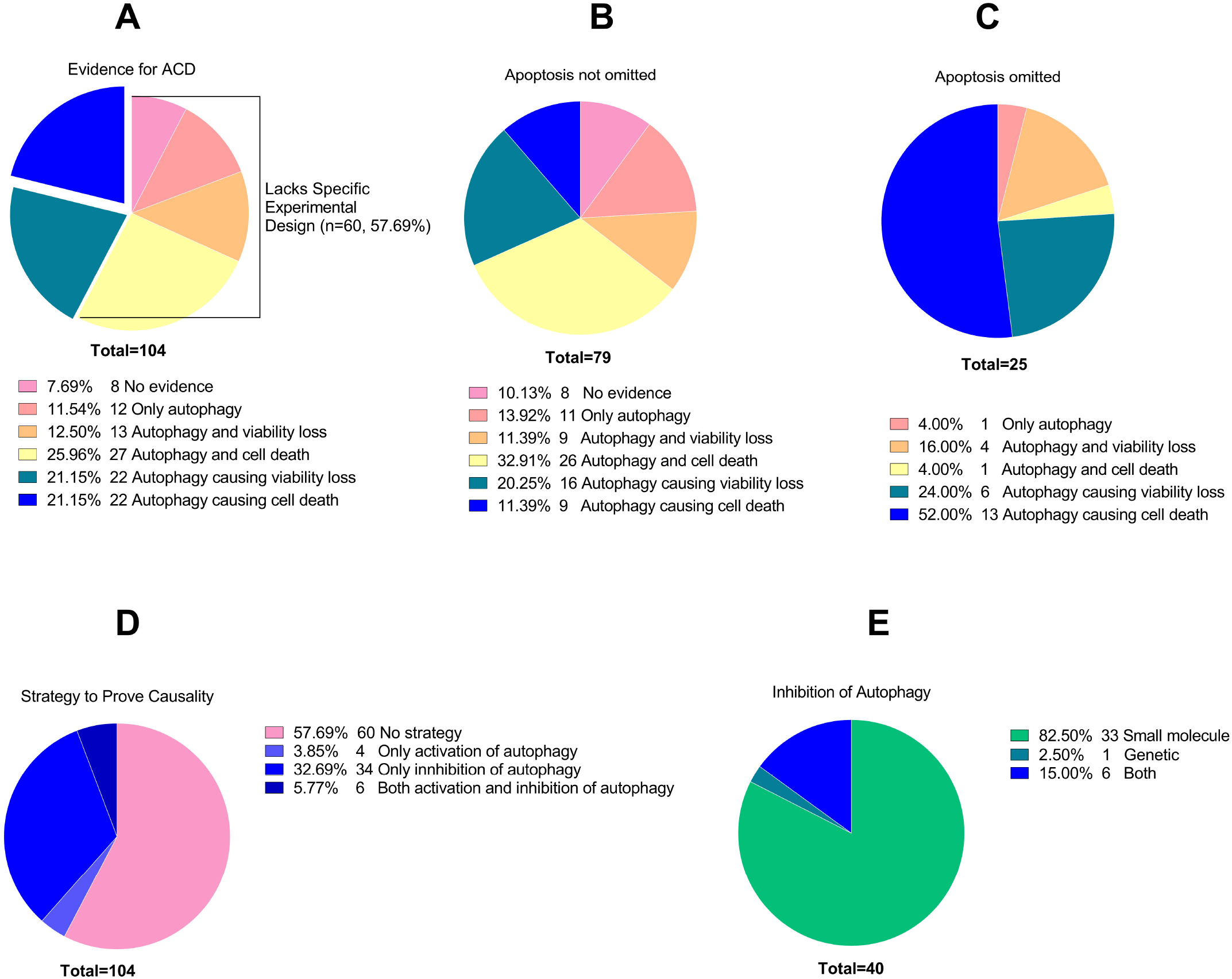
Distribution of articles based on evidence for autophagic cell death (ACD), strategies to prove causality, and types of autophagy inhibition. (A) Distribution of articles according to the evidence provided for ACD, without considering the status of apoptosis. (B) Distribution of articles that did not omit apoptosis as a potential mechanism of cell death. (C) Distribution of articles that omitted apoptosis to meet the criteria for ACD. (D) Distribution of articles based on the strategies employed to prove causality. (E) Distribution of articles based on the type of autophagy inhibition performed. In all graphs, numbers indicate the number of articles meeting the specified criteria.

In the remaining 40 articles (38.5%), there were experiments specifically designed to reveal the causal relationship between autophagy and cell death. However, since in 22 of these articles there was a reliance on viability loss instead of actual cell death measurement, only 22 remaining articles (21.2% of all articles) provided actual causality-level evidence for ACD (Figure 1A).

The most common strategy to prove causality was to inhibit autophagy and observe whether cell death or viability loss was prevented, a method used in 40 articles (Figure 1D). While 4 articles employed the opposite approach by activating autophagy and measuring the increase in cell death, in 6 articles (5.8%) both strategies are utilized to provide more robust evidence of causality (Figure 1D).

Of the 40 articles where autophagy was inhibited to prove causality, 33 relied solely on small molecule inhibitors, while 1 opted for genetic inhibition. 6 articles utilized both small-molecule and genetic-based inhibition to achieve more reliable results (Figure 1E). It is important to note that more articles used autophagy inhibitors overall. In total, small molecule inhibitors were used in 48, genetic inhibition was used in 5, both approaches together were used in 8 articles (see Supplementary Data). However, despite utilizing autophagy inhibition, 11 of these 61 articles did not measure viability and/or cell death following the inhibition, and thus failed to provide causality-level evidence.

Among all studies using small molecule inhibitors of autophagy, 3-methyladenine (3-MA) was the most popular choice, used in 27 studies, followed by chloroquine (CQ) in 26 studies and Bafilomycin A1 in 12 studies (see Supplementary Data). Four studies used non-specific inhibitors that are not listed in the ‘Genetic and Pharmacological Regulation of Autophagy’ table in the ‘Guidelines for the Use and Interpretation of Assays for Monitoring Autophagy’ [12]. Additionally, 13 studies employed more than one type of small molecule inhibitor, with 8 of them utilizing 3-MA and CQ (see Supplementary Data). In 18 of the studies, inhibition of autophagy was not demonstrated experimentally, thereby diminishing the robustness and reliability of their conclusions. Finally, only 27 out of 40 articles reported less than 50% rescue of cells from death after autophagy inhibition, suggesting that while autophagy may contribute to cell death, it may not the primary mechanism (See Supplementary Data).

### 5. Non-apoptotic evidence of cell death

Out of 104 articles, 14 (13.5%) failed to provide any evidence for cell death (Figure 2A). 36 articles (34.6%) demonstrated only viability loss and attributed that loss to cell death without further proof. Remaining 54 (51.92%) articles provided evidence for loss of membrane integrity as an indicator of cell death (Figure 2A).

**Figure 2.**
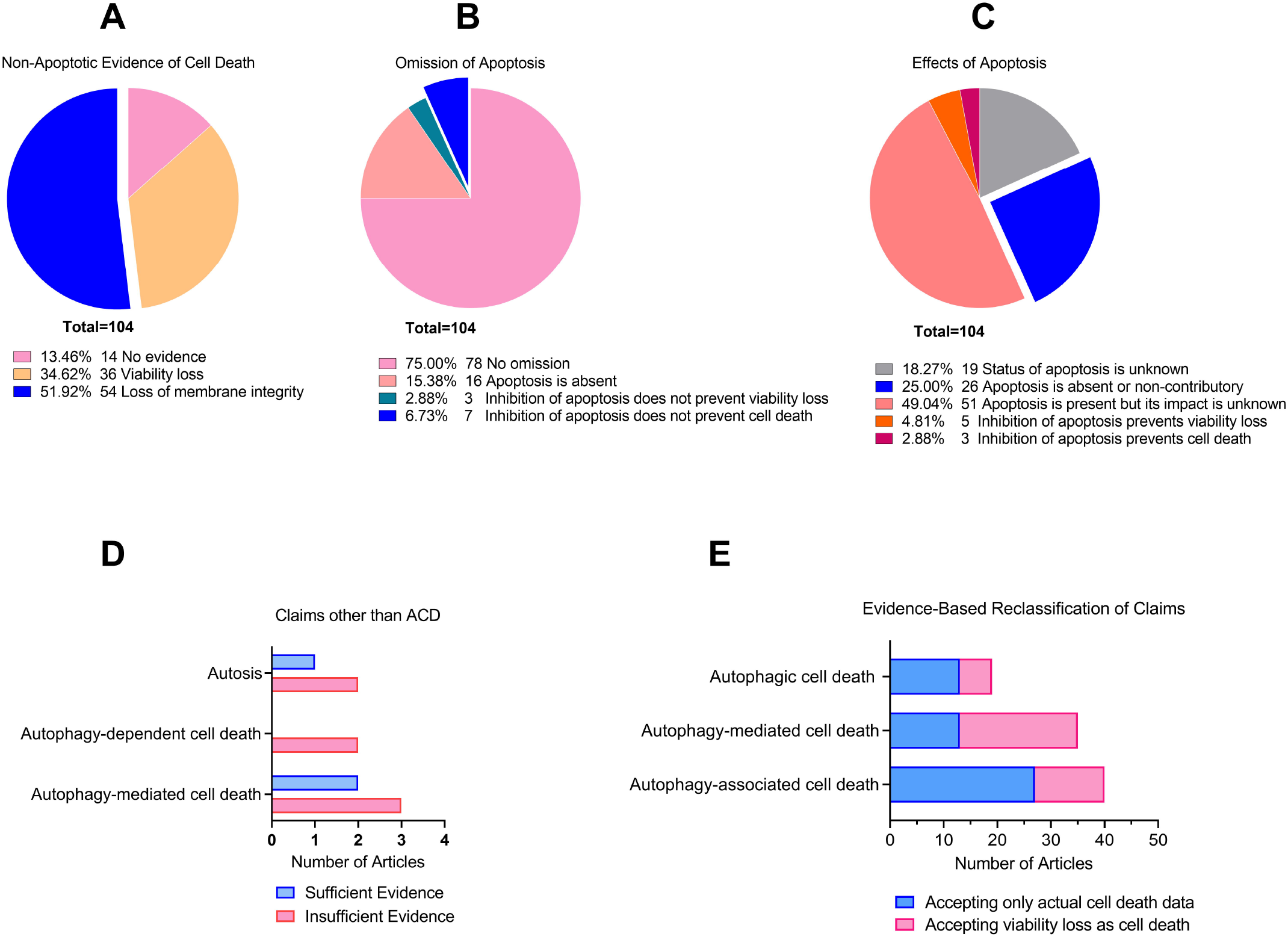
Distribution of articles based on evidence for cell death, strategies to omit apoptosis, effects of apoptosis, alternative claims, and evidence-based reclassification of claims. (A) Distribution of articles according to the evidence provided for cell death. (B) Distribution of articles based on the strategies employed to omit apoptosis. (C) Distribution of articles according to the cellular effects of apoptosis. (D) Distribution of articles based on claims other than ACD. Blue bars represent claims with sufficient evidence, while pink bars represent claims with insufficient evidence. (E) Distribution of articles reclassified based on the appropriate terminology assigned according to the evidence provided. Blue bars represent articles relying on cell death, and pink bars represent articles relying on viability loss. In all graphs, numbers indicate the number of articles meeting the specified criteria.

### 6. Experimental Design to Omit Apoptosis

Apoptosis was experimentally excluded in 26 articles (25.0%) (Figure 1C and 2B), with 16 of these articles simply demonstrating its absence. In contrast, 10 studies showed that apoptosis inhibitors were ineffective in preventing cell death and/or viability loss (Figure 2B).

Figure 1B presents data from studies that did not exclude apoptosis, while Figure 1C shows data from those that did. A comparison of these studies reveals that those omitting apoptosis (Figure 1C) had stronger experimental designs, with 52.00% demonstrating autophagy as a cause of cell death, compared to only 11.39% in studies that did not exclude apoptosis (Figure 1B).

### 7. Presence and Influence of Apoptosis, and Other Claims

Apoptotic status was unknown in 19 articles (19.27%) (Figure 2C). As mentioned earlier, 26 articles demonstrated that apoptosis either did not contribute to cell death or viability loss, or was completely absent.

In 8 articles, the inhibition of apoptosis prevented cell death and/or viability loss (Figure 2C), indicating that apoptosis was the primary mechanism of cell death. However, in 7 of these 8 studies, inhibition of autophagy also prevented cell death and/or viability loss (see Supplementary Data), suggesting that the mechanism should have been identified as autophagy-mediated apoptosis. While 2 of these articles accurately labelled the process, the remaining studies incorrectly referred to it as ‘autophagic cell death’.

Additionally, 3 other articles claimed ‘autophagy-mediated cell death’ but only demonstrated a correlation between autophagy and cell death, making ‘autophagy-associated cell death’ a more appropriate term in these cases (Supplementary data). In summary there were 5 articles claiming ‘autophagy-mediated cell death,’ but only 2 of them presented evidence that justified this classification.

Moreover, 11 articles demonstrated that inhibiting autophagy also inhibited apoptosis, a clear indication of ‘autophagy-mediated apoptosis’. However, all were incorrectly identified as ‘autophagic cell death’ by the authors. Similarly, 3 articles demonstrated that activating autophagy further intensified apoptosis which was again incorrectly identified as ‘autophagic cell death’ (see Supplementary Data).

2 articles claimed ‘autophagy-dependent cell death’, and both lacked the necessary supporting evidence. There were 3 articles claiming ‘autosis’, of which only 1 provided sufficient evidence for ACD (Figure 2D). Since the identification of ‘autophagy-dependent cell death’ and ‘autosis’ also requires satisfying the established criteria for ACD, we analysed these claims accordingly.

### 8. Reclassification of Autophagy-Related Cell Death Claims Based on Evidence Provided

The graph (Figure 2E) illustrates the distribution of articles according to the appropriate terminology assigned based on the evidence they provided, rather than the claims explicitly made by the authors. Specifically, it highlights the number of articles that should have been classified under the terms ‘autophagic cell death’, ‘autophagy-mediated cell death’, or ‘autophagy-associated cell death’, based on their experimental evidence. Specifically, ACD claims were accepted only when a causal relationship between autophagy and cell death was demonstrated, alongside the omission of apoptosis, as recommended by established guidelines [6], [8], [9], [12]. This data can also be seen in Figure 1C. The graph further distinguishes between studies relying on cell death evidence (blue bars) and viability loss data (pink bars). This differentiation underscores the significant reliance on less stringent criteria by nearly half of the studies, as viability loss does not necessarily equate to direct cell death. It is important to underline that since we only included studies with a type of ‘autophagic cell death’ claim, in ideal conditions the number of articles re-classified as ‘autophagy-associated cell death’ should have been zero. It is also important to note that 21 studies provided no evidence for any type of claim and are therefore not represented in the graph (see Supplementary data).

## Discussion

Autophagy is a crucial mechanism with implications for a wide range of physiological and pathological processes, including cancer, neurodegenerative diseases, and immune responses. Despite its importance, the precise nature of autophagy as a cell death mechanism remains a topic of intense debate, making the accurate identification of autophagic cell death (ACD) essential for advancing research in this field.

In this study, we systematically evaluated 104 research articles that claimed to demonstrate ACD, uncovering significant limitations in the experimental designs utilized. While less than half of the articles presented evidence suggesting a causal relationship between autophagy and cell death, fifteen of these relied on viability loss rather than direct measurements of cell death, leaving only 22 (21.2%) studies that demonstrated autophagy as a cause of cell death. However, according to established guidelines, the exclusion of apoptosis is also a crucial criterion for accurately identifying ACD [6], [8], [9], [12]. When this criterion was applied, only 13 articles (12.5%) met all the necessary requirements, highlighting a substantial gap between claimed and verified evidence.

However, a counter-argument could suggest that the authors may have used the term ‘ACD’ as a shorthand to indicate the correlation between autophagy and cell death, even if such a practice is not recommended and the term ‘autophagy-associated cell death’ should have been used instead [6], [12]. Nonetheless, even if this were the case, our analysis revealed 8 articles with no evidence whatsoever, 11 articles that demonstrated only autophagy without providing any information on cell death, and an additional 9 articles that relied solely on viability loss rather than direct evidence of cell death.

Notably, the previously proposed nomenclature does not appear to be widely adopted by researchers. The vast majority of claims referred to ACD, while the usage of other terms was quite rare. For instance, only four articles (one of which was omitted as it did not include autophagic cell death claim) used the term ‘autophagy-associated cell death’ but in reality, more studies demonstrated ‘autophagy-associated cell death,’ which requires only correlation-level evidence. There were a handful of other claims including five ‘autophagy-mediated’, three ‘autosis’, and two ‘autophagy-dependent’ cell deaths, with only four articles providing the necessary evidence (two for ‘autophagy-mediated’ and one for ‘autosis’). When examining the actual evidence provided by the studies, 27 demonstrated ‘autophagy-associated cell death’, 13 identified ‘autophagy-mediated cell death’, and 13 provided evidence for ‘autophagic cell death’ (Figure 2E).

In addition to challenges with nomenclature, our findings highlight concerning practices, which further complicate the interpretation of results, including the reliance on viability data instead of direct measurement of cell death, which was observed in 34 studies overall (32.7%) and in 22 studies which provided causality-level evidence (50.0%).

Other concerning practices were observed in studies investigating autophagy and its inhibition. Notably, only 47 studies (45.19%) demonstrated the full autophagic process (autophagic flux) from autophagosome formation to lysosomal degradation. The remaining studies relied solely on static markers, which do not necessarily indicate active degradation and may lead to misinterpretation of autophagy status [16]. However, since demonstrating both autophagic flux and a causal relationship between autophagy and cell death requires the use of autophagy inhibitors, all 13 articles that met the full criteria for autophagic cell death also provided evidence of autophagic flux.

The most commonly used inhibitor was 3-methyladenine (3-MA), a non-specific phosphatidylinositol 3-phosphate inhibitor that blocks the early stages of autophagy [12] but can also promote autophagy in certain systems [17], and it was employed in 27 of 40 studies (67.5%). Practices to enhance robustness, such as using multiple inhibitors (13 of 40 studies, 32.5%) or combining genetic inhibition with small-molecule inhibitors (6 of 40 studies, 15.0%), were relatively uncommon. Additionally, 18 of 40 studies (45.0%) utilized autophagy inhibitors like 3-MA or CQ without experimentally demonstrating the inhibition of autophagy. In cases where inhibition evidence was provided, it often involved an increase in LC3-II levels. While this is an acceptable indicator of flux inhibition when using agents such as Bafilomycin A1 or CQ, both of which block LC3-II turnover [12], its simultaneous use as a marker of autophagy induction within the same studies complicates the interpretation of autophagy inhibition.

Similarly, apoptosis inhibitors, such as caspase inhibitors, were often used without verifying that apoptosis was inhibited. In four studies, apoptosis was not measured at any point, neither before nor after inhibitor treatment, despite the use of apoptotic inhibitors (see Supplementary file).

In summary, our analysis reveals significant inconsistencies and limitations in the current literature on autophagic cell death, particularly concerning experimental rigor and terminological precision. Only a small fraction of studies (12.5%) met established criteria for ACD, underscoring the need for more robust experimental designs and adherence to standardized guidelines. To aid researchers in addressing these shortcomings, we propose a systematic workflow (Figure 3) that outlines the experimental and analytical steps required to classify evidence for different modes of autophagy accurately, also addressing the issue of relying on viability loss as an indicator of cell death. This flowchart provides a framework for obtaining correlation-or causality-level evidence, emphasizing key decision points such as the exclusion of apoptosis and the role of autophagy inhibition. Addressing these shortcomings will not only advance our understanding of the role of autophagy in cell death but also reflects the broader responsibility of the scientific community, including authors and reviewers, to uphold rigor and precision in both research and its evaluation.

**Figure 3.**
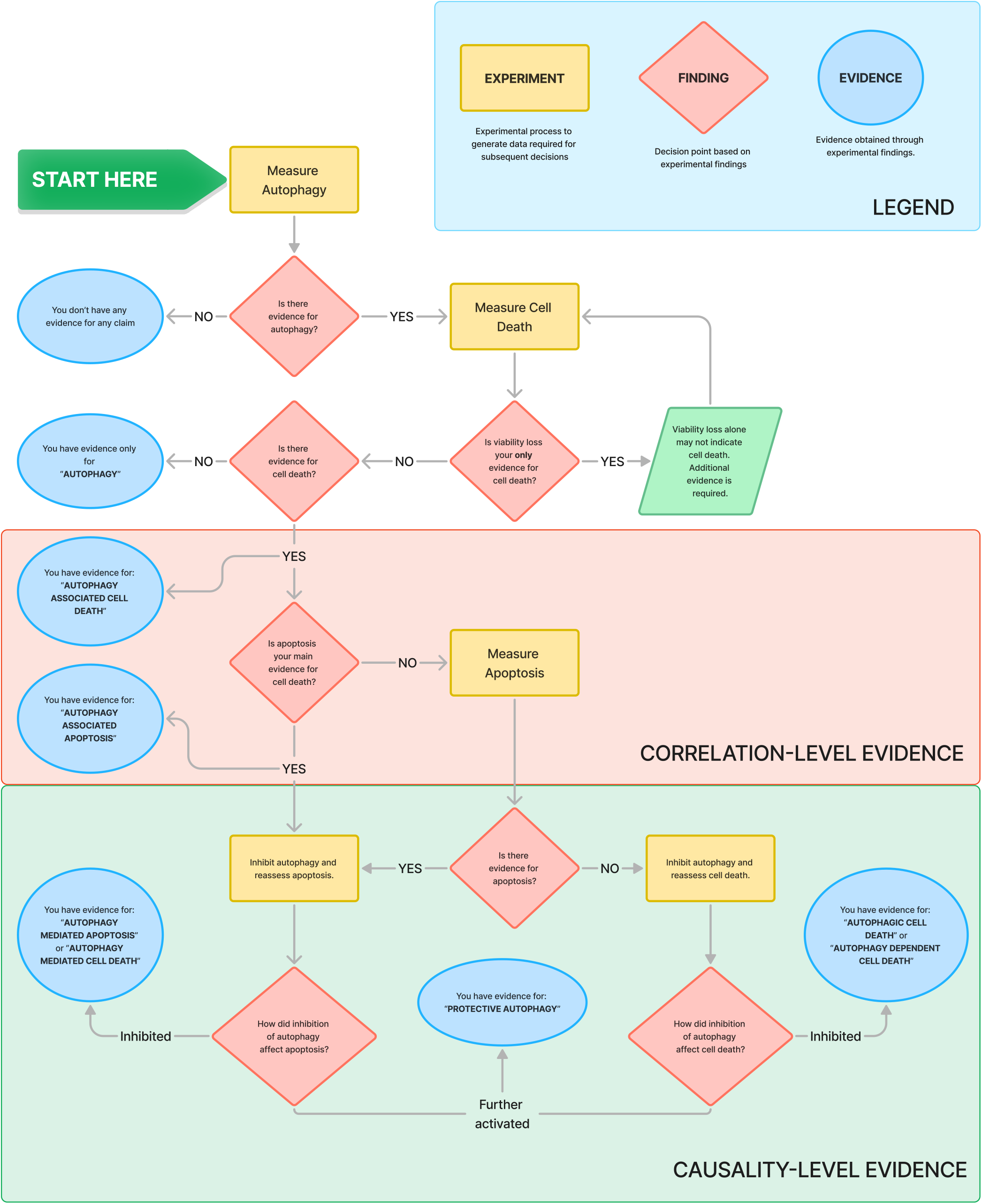
Flowchart illustrating the systematic approach to classify evidence for identifying different modes of autophagy. The flowchart guides researchers through experimental steps and decision points to obtain evidence for various types of autophagy, including ‘autophagy-associated cell death,’ ‘autophagy-mediated cell death,’ ‘autophagy-dependent cell death,’ and ‘protective autophagy.’

## Abbreviations

ACD: Autophagic Cell Death

## Author Contributions

ABÖ conceived and designed the study, determined the methodology, supervised data analysis, and wrote the manuscript. YG conducted the main data collection and analysis and contributed to manuscript writing. Both authors reviewed and approved the final version of the manuscript.

## Acknowledgements

We would like to thank Kıvanç Görgülü for his assistance in accessing the full texts of certain articles.

## Data Availability Statement

The data that supports the findings of this study are available in figure 1, figure 2, and the supplementary material of this article.

## Supporting Information

**Data S1**. Database of all 104 articles with ACD claims and evidence (15 fields).

