## Supplementary Document for "Most autophagic cell death studies lack evidence of causality"

Supplementary Table. The table contains the complete database of all evaluated articles, with each row representing an individual article. Columns detail the study methodologies and experimental designs, providing a comprehensive analysis of the evidence and criteria assessed. There are 15 columns in total, as follows:

1. Article Name
2. ID - A unique number for ease of referencing
3. Article DOI Number
4. Journal Name
5. Is there a claim of ACD? – Presence or absence of an autophagic cell death (ACD) claim
6. What is the claim? – Type of the claim (ACD, Autophagy Associated Cell Death, etc.)
7. Evidence of Autophagy – How autophagy was identified (LC3II, decreased P62, autophagosomes, etc.)
8. Experimental Model – including the names of the cell lines or animal models used in the study
9. Interference to Induce ACD – including small molecules, siRNA or gene overexpression
10. Evidence for Positive Autophagic Flux – including LC3 turnover, p62 degradation, or autophagolysosome formation.
11. Autophagy Inhibition Type – including genetic, small molecule, or both (not performed if there is no inhibition)
12. Evidence of Cell Death (Except for Apoptosis) – including reduced metabolic activity, loss of membrane integrity, etc.
13. Evidence of Apoptosis – including caspase activity, phosphatidyl serine exposure, DNA fragmentation, etc.
14. Status of Apoptosis – Whether apoptosis was excluded, present, or not addressed.
15. Evidence of ACD – experimental approach embraced to prove ACD.

The “Status of Apoptosis” and “Evidence of ACD” columns use symbolic shorthand: a plus sign before a process (e.g., +Autophagy) indicates induction, while a minus sign before a process (e.g., -Apoptosis) indicates inhibition. A plus or minus sign after a process (e.g., Cell Death+, Apoptosis-) reflects an increase or decrease in that outcome. The greater-than symbol (e.g., +Autophagy > Cell Death+) indicates a causal relationship. “X” means no change was observed, and “unknown” indicates insufficient information to determine the status of that particular process. This symbolic notation is further explained by the examples provided below;

- Example 1: Autophagy+ means an increase in autophagy
- Example 2: Viability- means a decrease in cell viability
- Example 3: ApoptosisX means no change in apoptosis
- Example 4: +Autophagy > Cell Death+ means induction of autophagy leads to an increase in cell death
- Example 5: -Autophagy > Cell Death- means inhibition of autophagy leads to a decrease in cell death
- Example 6: -Apoptosis >Autophagy X means inhibition of apoptosis leads to no change in autophagy

| **Article Name** | **ID** | **Article DOI** | **Journal Name** | **Is there a claim of ACD?** | **What is the claim?** | **Evidence of Autophagy** | **Experimental Model** | **Interference to Induce ACD** | **Evidence for Positive Autophagic Flux** | **Autophagy Inhibition Type** | **Evidence of Cell Death (Except for Apoptosis)** | **Evidence of Apoptosis** | **Status of Apoptosis** | **Evidence of ACD** |
| --- | --- | --- | --- | --- | --- | --- | --- | --- | --- | --- | --- | --- | --- | --- |
| Pd(II) and Rh(III) Complexes with Isoquinoline D  erivatives Induced Mitochondria-Mediated Apoptotic and Autophagic Cell Death in HepG2 Cells | 1 | 10.31635/ccschem.020.202000363 | CCS CHEMISTRY | Yes | Autophagic Cell Death (ACD) | LC3-II, p62 | Cell line, human hepatocellular carcinoma (HepG2) | Small Molecule Inhibitor | No evidence | Not performed | Phosphatidyl serine exposure/PI internalization | Phosphatidyl serine exposure, DNA fragmantation | Apoptosis+ | Autophagy+  Cell Death+ |
| Mechanisms Involved in Ceramide-Induced Autophagy in Osteoblasts | 2 | 10.51847/B8jiA53A5q | ARCHIVES OF PHARMACY PRACTICE | Yes | Autophagic Cell Death (ACD) | LC3-II, Visualization of autophagosomes | Cell line, murine osteogenic (CHME3) | Small Molecule Inhibitor | Autophagolysosome Formation , LC3-II Turnover | Genetic | Loss of membrane integrity | Nuclear margination/ Chromatin condensation, Membrane blebbing | Apoptosis+ | Autophagy+  Cell Death+ |
| Electrospun meshes intrinsically promote M2 polarization of microglia under hypoxia and offer protection from hypoxia-driven cell death | 3 | 10.1088/1748-605X/ac0a91 | BIOMEDICAL MATERIALS | Yes | Autophagic Cell Death (ACD) | Not available | Cell line, human microglial (CHME3) | Small Molecule Inhibitor | No evidence | Not performed | Reduced metabolic activity | Caspase activation | Apoptosis+ | None  Viability- |
| DUSP4 inhibits autophagic cell death in PTC by inhibiting JNK-BCL2-Beclin1 signaling | 4 | 10.1139/bcb-2020-0636 | BIOCHEMISTRY AND CELL BIOLOGY | Yes | Autophagic Cell Death (ACD) | LC3-II | Cell line, human papillary thyroid carcinoma cell line (K1), human papillary thyroid carcinoma cell line (TPC-1) | RNA Interference | No evidence | Not performed | Loss of membrane integrity, Phosphatidyl serine exposure/PI internalization | DNA fragmantation, Phosphatidyl serine exposure | Apoptosis+ | Autophagy+  Cell Death+ |
| Qing Yan Li Ge Tang, a Chinese Herbal Formula, Induces Autophagic Cell Death through the PI3K/Akt/mTOR Pathway in Nasopharyngeal Carcinoma Cells In Vitro | 5 | 10.1155/2021/9925684 | EVIDENCE-BASED COMPLEMENTARY AND ALTERNATIVE MEDICINE | Yes | Autophagic Cell Death (ACD) | Not available | Cell line, nasopharyngeal carcinoma cell line (NPC-TW01), nasopharyngeal carcinoma cell line (HONE-1) | Small Molecule Inhibitor | No evidence | Small Molecule, 3MA | Reduced colony formation, Reduced metabolic activity | Caspase activation | ApoptosisX | Viability-  None |
| Autophagy Prevents Osteocyte Cell Death under Hypoxic Conditions | 6 | 10.1159/000519086 | CELLS TISSUES ORGANS | No |  | Not available | Cell line, murine osteocyte-like cell line (MLO-Y4) |  |  | Small Molecule, 3MA | None | None | Unknown | None |
| Apigenin Induces Autophagy and Cell Death by Targeting EZH2 under Hypoxia Conditions in Gastric Cancer Cells | 7 | 10.3390/ijms222413455 | INTERNATIONAL JOURNAL OF MOLECULAR SCIENCES | Yes | Autophagic Cell Death (ACD) | LC3-II, p62 | Cell line, "human gastric cancer cell lines (AGS, SNU-216, NCI-N87, SNU-638, MKN-7, and MKN-74)" | Small Molecule Inhibitor | LC3-II Turnover , p62 Degradation and Rescue | Both, 3MA, CQ | Loss of membrane integrity | Caspase activation | -Apoptosis > Cell Death- | -Autophagy > Cell Death-  +Autophagy > Cell Death+ |
| Repurposing Antipsychotics for Cancer Treatment | 8 | 10.3390/biomedicines9121785 | BIOMEDICINES | No |  |  |  |  |  |  |  |  |  |  |
| AMPK-Mediated Metabolic Switching Is High Effective for Phytochemical Levo-Tetrahydropalmatine (l-THP) to Reduce Hepatocellular Carcinoma Tumor Growth | 9 | 10.3390/metabo11120811 | METABOLITES | No |  | LC3-II |  |  |  | Not performed | Reduced metabolic activity |  |  | Autophagy+  Viability- |
| Discovery of Potent and Novel Dual PARP/BRD4 Inhibitors for Efficient Treatment of Pancreatic Cancer | 10 | 10.1021/acs.jmedchem.1c01535 | JOURNAL OF MEDICINAL CHEMISTRY | No | Autophagy Associated Cell Death (AACD) | p62 |  |  |  | Not performed | Phosphatidyl serine exposure/PI internalization | Phosphatidyl serine exposure |  | Autophagy+  Cell Death+ |
| Snake venom induces an autophagic cell death via activation of the JNK pathway in colorectal cancer cells | 11 | 10.7150/jca.75791 | JOURNAL OF CANCER | Yes | Autophagic Cell Death (ACD) | LC3-II, p62 | Cell line, human colon adenocarcinoma cell line (SW480), Human lung adenocarcinoma cell line (A549), human cervical cancer cell line (HeLa), human osteosarcoma cell line (U-2 OS), human normal colon cell line (CCD-18Co) | Small Molecule Inhibitor | LC3-II Turnover , p62 Degradation and Rescue | Small Molecule, CQ, Non specific Small Molecule, JNK inhibitor | Loss of membrane integrity, Reduced metabolic activity | DNA fragmantation | Apoptosis+ | -Autophagy > Cell Death- |
| Aloin promotes oral squamous cell carcinoma cell apoptosis and autophagy through Akt/mTOR pathway | 12 | 10.15586/qas.v14i2.978 | QUALITY ASSURANCE AND SAFETY OF CROPS & FOODS | No |  | LC3-II, p62 |  |  |  | Small Molecule, CQ |  | Caspase activation |  | Autophagy+  Cell Death+ |
| THE DIARYL-IMIDAZOPYRIDAZINE ANTI-PLASMODIAL COMPOUND, MMV652103, EXHIBITS ANTI-BREAST CANCER ACTIVITY | 13 | 10.17179/excli2021-4323 | EXCLI JOURNAL | Yes | Autophagic Cell Death (ACD) | LC3-II | Cell line, human breast cancer cell line (MCF7), human breast cancer cell line (T47D), human breast cancer cell line (MDA-MB-468), human breast cancer cell line (MDA-MB-231), human breast cancer cell line (BT474) | Small Molecule Inhibitor | No evidence | Genetic | None | DNA fragmantation, Cleaved PARP | -Autophagy > Apoptosis-Apoptosis+ | Autophagy+  Cell Death+ |
| Falcarindiol Stimulates Apoptotic and Autophagic Cell Death to Attenuate Cell Proliferation, Cell Division, and Metastasis through the PI3K/AKT/mTOR/p70S6K Pathway in Human Oral Squamous Cell Carcinomas | 14 | 10.1142/S0192415X22500112 | AMERICAN JOURNAL OF CHINESE MEDICINE | Yes | Autophagic Cell Death (ACD) | LC3-II, Visualization of autophagosomes | Cell line, human oral squamous cell carcinoma cell line (YD-10B) | Small Molecule Inhibitor | No evidence | Not performed | Reduced metabolic activity | DNA fragmantation, Caspase activation | Apoptosis+ | Autophagy+  Cell Death+ |
| Cell death mechanisms induced by synergistic effects of halofuginone and artemisinin in colorectal cancer cells | 15 | 10.7150/ijms.66737 | INTERNATIONAL JOURNAL OF MEDICAL SCIENCES | Yes | Autophagic Cell Death (ACD) | LC3-II, p62 | Cell line, human colorectal cancer cell line (DLD-1), human colorectal cancer cell line (HCT116) | Small Molecule Inhibitor | LC3-II Turnover , p62 Degradation and Rescue | Small Molecule, CQ | Loss of membrane integrity | Caspase activation | -Apoptosis > Autophagy+  -Apoptosis > Cell Deathx | +Autophagy > Cell Death+ |
| Tfeb-Mediated Transcriptional Regulation of Autophagy Induces Autosis during Ischemia/Reperfusion in the Heart | 16 | 10.3390/cells11020258 | CELLS | Yes | Autosis | Visualization of autophagosomes | Animal Tissue, Animal Primary Cells, C57BL/6J mouse hearts, Primary neonatal rat cardiomyocytes (NRCMs) | Gene Over-expression | Autophagolysosome Formation | Genetic | Reduced metabolic activity | None | Unknown | Viability-  Autophagy+ |
| Plumbagin triggers redox-mediated autophagy through the LC3B protein in human papillomavirus-positive cervical cancer cells | 17 | 10.5114/aoms.2020.101072 | ARCHIVES OF MEDICAL SCIENCE | Yes | Autophagic Cell Death (ACD) | LC3-II, Visualization of autophagosomes | Cell line, human cervical cancer cell line (SiHa) | Small Molecule Inhibitor | Autophagolysosome Formation | Not performed | Loss of membrane integrity | DNA fragmantation, Membrane blebbing | Apoptosis+ | Autophagy+  Cell Death+ |
| Suppression of DYRK1A/B Drives Endoplasmic Reticulum Stress-mediated Autophagic Cell Death Through Metabolic Reprogramming in Colorectal Cancer Cells | 18 | 10.21873/anticanres.15516 | ANTICANCER RESEARCH | No |  |  |  |  |  |  |  |  |  |  |
| Magnesium Isoglycyrrhizinate Ameliorates Concanavalin A-Induced Liver Injury by Inhibiting Autophagy | 19 | 10.3389/fphar.2021.794319 | FRONTIERS IN PHARMACOLOGY | Yes | Autophagic Cell Death (ACD) | LC3-II, p62 | Animal Tissue, Animal Primary Cells, Liver tissues from Balb/c mice, Primary mice hepatocytes | Small Molecule Inhibitor | LC3-II Turnover , p62 Degradation and Rescue | Small Molecule | Loss of membrane integrity | Caspase activation, DNA fragmantation | ApoptosisX  +Autophagy >Apoptosis+ | +Autophagy > Cell Death+ |
| Biotinylated curcumin as a novel chemosensitizer enhances naphthalimide-induced autophagic cell death in breast cancer cells | 20 | 10.1016/j.ejmech.2021.114029 | EUROPEAN JOURNAL OF MEDICINAL CHEMISTRY | Yes | Autophagic Cell Death (ACD) | LC3-II, p62 | Cell line, human breast cancer cell line (MCF7), mouse skin melanoma cell line (B16), normal human lung fibroblast cell line (HLF) | Small Molecule Inhibitor | No evidence | Not performed | Loss of membrane integrity, Reduced metabolic activity | Phosphatidyl serine exposure | ApoptosisX | Autophagy+  Cell Death+ |
| S-20, a steroidal saponin from the berries of black nightshade, exerts anti-multidrug resistance activity in K562/ADR cells through autophagic cell death and ERK activation | 21 | 10.1039/d1fo03191k | FOOD & FUNCTION | Yes | Autophagic Cell Death (ACD) | LC3-II, p62 | Cell line, Human chronic myelogenous leukemia cell line (K562), Human adriamycin-resistant chronic myelogenous leukemia cell line (K562/ADR), Human promyelocytic leukemia cell line (HL-60), Human histiocytic lymphoma cell line (U937), Human T-cell leukemia cell line (Jurkat) | Small Molecule Inhibitor | LC3-II Turnover , p62 Degradation and Rescue | Small Molecule, 3MA, CQ | Reduced metabolic activity | Caspase activation, Cleaved PARP, Phosphatidyl serine exposure | -Apoptosis >Autophagy X  -Apoptosis > Viability+  -Autophagy > Apoptosis- | -Autophagy > Viability+ |
| Deferoxamine Counteracts Cisplatin Resistance in A549 Lung Adenocarcinoma Cells by Increasing Vulnerability to Glutamine Deprivation-Induced Cell Death | 22 | 10.3389/fonc.2021.794735 | FRONTIERS IN ONCOLOGY | Yes | Autophagic Cell Death (ACD) | LC3-II | Cell line, Human lung adenocarcinoma cell line (A549), Human cisplatin-resistant lung adenocarcinoma cell line (A549/DDP) | Small Molecule Inhibitor | LC3-II Turnover | Small Molecule, JNK inhibitor, Bafilomycin | Phosphatidyl serine exposure/PI internalization | Caspase activation, Phosphatidyl serine exposure | -Apoptosis > Viability+ | Cell Death+  Autophagy+ |
| Development of a model to predict the prognosis of esophageal carcinoma based on autophagy-related genes | 23 | 10.2217/fon-2021-0070 | FUTURE ONCOLOGY | No |  |  |  |  |  |  |  |  |  |  |
| ACT001 inhibits pituitary tumor growth by inducing autophagic cell death via MEK4/MAPK pathway | 24 | 10.1038/s41401-021-00856-5 | ACTA PHARMACOLOGICA SINICA | Yes | Autophagic Cell Death (ACD) | LC3-II | Cell line, Rat pituitary tumor cell line (GH3), Rat pituitary tumor cell line (MMQ) | Small Molecule Inhibitor | LC3-II Turnover | Both, 3MA | Phosphatidyl serine exposure/PI internalization | Phosphatidyl serine exposure | Apoptosis+ | -Autophagy > Viability+ |
| Autophagic and apoptotic cell death induced by the quinoline derivative 2-(6-methoxynaphthalen-2-yl)quinolin-4-amine in pancreatic cancer cells is via ER stress and inhibition of Akt/mTOR signaling pathway | 25 | 10.1002/ddr.21916 | DRUG DEVELOPMENT RESEARCH | Yes | Autophagic Cell Death (ACD) | LC3-II, p62 | Cell line, Human pancreatic cancer cell line (PANC-1), Human pancreatic cancer cell line (MIA PaCa-2), Chinese hamster ovary cell line (CHO) | Small Molecule Inhibitor | LC3-II Turnover , p62 Degradation and Rescue | Small Molecule, CQ | None | Caspase activation, Cleaved PARP | -Autophagy > Apoptosis--Apoptosis > Viability+ | -Autophagy > Viability+ |
| Main active components of Si-Miao-Yong-An decoction (SMYAD) attenuate autophagy and apoptosis via the PDE5A-AKT and TLR4-NOX4 pathways in isoproterenol (ISO)-induced heart failure models | 26 | 10.1016/j.phrs.2022.106077 | PHARMACOLOGICAL RESEARCH | Yes | Autophagic Cell Death (ACD) | LC3-II, p62 | Cell line, Animal Tissue, Rat cardiomyoblast cell line (H9c2), Rat cardiac tissue | Small Molecule Inhibitor | LC3-II Turnover , p62 Degradation and Rescue | Small Molecule, 3MA, CQ | Phosphatidyl serine exposure/PI internalization | Caspase activation, Phosphatidyl serine exposure | -Autophagy > Apoptosis  +Autophagy >Apoptosis+  Apoptosis+ | Cell Death+  Autophagy+ |
| A Novel Algicidal Bacterium, Microbulbifer sp. YX04, Triggered Oxidative Damage and Autophagic Cell Death in Phaeocystis globosa, Which Causes Harmful Algal Blooms | 27 | 10.1128/spectrum.00934-21 | MICROBIOLOGY SPECTRUM | Yes | Autophagic Cell Death (ACD) | LC3-II | Cell line, Phaeocystis globosa (CCMA-191) | Small Molecule Inhibitor | No evidence | Not performed | Loss of membrane integrity | None | Unknown | Autophagy+  Cell Death+ |
| CD142 plays a key role in the carcinogenesis of gastric adenocarcinoma by inhibiting BCL2-dependent autophagy | 28 | 10.1139/bcb-2021-0144 | BIOCHEMISTRY AND CELL BIOLOGY | Yes | Autophagic Cell Death (ACD) | LC3-II, Visualization of autophagosomes | Cell line, Human normal gastric epithelial cell line (GES-1), Human gastric adenocarcinoma cell line (SNU16), Human gastric adenocarcinoma cell line (SGC083), Human gastric adenocarcinoma cell line (MKN45), Human gastric adenocarcinoma cell line (HGC27) | RNA Interference | LC3-II Turnover | Small Molecule, 3MA | Phosphatidyl serine exposure/PI internalization, Loss of membrane integrity | Phosphatidyl serine exposure | Apoptosis+ | Cell Death+  Autophagy+ |
| Alterations of sodium-hydrogen exchanger 1 function in response to SGLT2 inhibitors: what is the evidence? | 29 | 10.1007/s10741-022-10220-2 | HEART FAILURE REVIEWS | No |  |  |  |  |  |  |  |  |  |  |
| Enhancement of anti-PD-1/PD-L1 immunotherapy for osteosarcoma using an intelligent autophagy-controlling metal organic framework | 30 | 10.1016/j.biomaterials.2022.121407 | BIOMATERIALS | Yes | Autophagic Cell Death (ACD) | LC3-II, p62 | Cell line, Mouse osteosarcoma cell line (K7M2), Mouse pre-osteoblast cell line (MC3T3-E1) | Small Molecule Inhibitor | LC3-II Turnover , p62 Degradation and Rescue | Both, 3MA | Loss of membrane integrity | None | Unknown | -Autophagy > Cell Death- |
| Targeting ferroptosis in acute kidney injury | 31 | 10.1038/s41419-022-04628-9 | CELL DEATH & DISEASE | No |  |  |  |  |  |  |  |  |  |  |
| Disulfiram/copper induces antitumor activity against gastric cancer cells in vitro and in vivo by inhibiting S6K1 and c-Myc | 32 | 10.1007/s00280-022-04398-3 | CANCER CHEMOTHERAPY AND PHARMACOLOGY | Yes | Autophagic Cell Death (ACD) | LC3-II | Cell line, Human gastric adenocarcinoma cell line (HGC27), Human gastric cancer cell line (SGC-7901) | Small Molecule Inhibitor | No evidence | Not performed | Phosphatidyl serine exposure/PI internalization | Caspase activation, Phosphatidyl serine exposure | Apoptosis+ | Autophagy+ |
| Leukemia inhibitory factor prevents chicken follicular atresia through PI3K/AKT and Stat3 signaling pathways | 33 | 10.1016/j.mce.2021.111550 | MOLECULAR AND CELLULAR ENDOCRINOLOGY | Yes | Autophagic Cell Death (ACD) | LC3-II, p62 | Animal Primary Cells, Primary granulosa cells (GCs), Animal Tissue, Chicken ovarian follicle tissue | Small Molecule Inhibitor | LC3-II Turnover , p62 Degradation and Rescue | Small Molecule, PI3K inhibitor | None | DNA fragmantation, Caspase activation, Nuclear margination/ Chromatin condensation | +Autophagy > Apoptosis+  Apoptosis+ | +Autophagy > Viability- |
| Hexadecanoic acid-enriched extract of Halymenia durvillei induces apoptotic and autophagic death of human triple-negative breast cancer cells by upregulating ER stress | 34 | 10.4103/2221-1691.338922 | ASIAN PACIFIC JOURNAL OF TROPICAL BIOMEDICINE | Yes | Autophagic Cell Death (ACD) | LC3-II | Cell line, human breast cancer cell line (MDA-MB-231) | Small Molecule Inhibitor | No evidence | Not performed | Phosphatidyl serine exposure/PI internalization, Reduced metabolic activity | Caspase activation, Phosphatidyl serine exposure | Apoptosis+ | Autophagy+  Cell Death+ |
| FGA inhibits metastases and induces autophagic cell death in gastric cancer via inhibiting ITGA5 to regulate the FAK/ERK pathway | 35 | 10.1016/j.tice.2022.101767 | TISSUE & CELL | Yes | Autophagic Cell Death (ACD) | LC3-II, p62 | Cell line, Human gastric cancer cell line (NCI-N87), Human gastric adenocarcinoma cell line (HGC27), Human gastric cancer cell line (HS-746T), Human normal gastric epithelial cell line (GES-1) | Gene Over-expression | No evidence | Not performed | Reduced metabolic activity | None | Unknown | Autophagy+  Viability- |
| NAD plus depletion enhances reovirus-induced oncolysis in multiple myeloma | 36 | 10.1016/j.omto.2022.02.017 | MOLECULAR THERAPY-ONCOLYTICS | Yes | Autophagic Cell Death (ACD) | LC3-II, p62 | Cell line, "Human multiple myeloma cell lines (KMS11, IM9, MM1S, U226, JJN3, KMM1, KMS12PE, MOLP8, RPMI8226, KMS27, MM1R)" | Small Molecule Inhibitor | LC3-II Turnover , p62 Degradation and Rescue | Small Molecule, CQ | Loss of membrane integrity, Phosphatidyl serine exposure/PI internalization | Phosphatidyl serine exposure | Apoptosis+ | -Autophagy > Cell Death- |
| Characterization of KRC-108 as a TrkA Kinase Inhibitor with Anti-Tumor Effects | 37 | 10.4062/biomolther.2021.195 | BIOMOLECULES & THERAPEUTICS | Yes | Autophagy Mediated Cell Death | LC3-II, p62 | Cell line, Human colon cancer cell line (KM12C) | Small Molecule Inhibitor | No evidence | Not performed | Phosphatidyl serine exposure/PI internalization | Phosphatidyl serine exposure | Apoptosis+ | Cell Death+  Autophagy+ |
| alpha-Hederin inhibits the platelet activating factor-induced metastasis of HCC cells through disruption of PAF/PTAFR axis cascaded STAT3/MP-2 expression | 38 | 10.1016/j.phrs.2022.106180 | PHARMACOLOGICAL RESEARCH | No |  | LC3-II |  |  |  | Genetic | Loss of membrane integrity | Nuclear margination/ Chromatin condensation, Membrane blebbing |  |  |
| F-box protein FBXO41 plays vital role in arsenic trioxide-mediated autophagic death of cancer cells | 39 | 10.1016/j.taap.2022.115973 | TOXICOLOGY AND APPLIED PHARMACOLOGY | Yes | Autophagic Cell Death (ACD) | p62, LC3-II | Cell line, human breast cancer cell line (MCF7), Human non-tumorigenic breast epithelial cell line (MCF10A), Human embryonic kidney cell line (HEK-293T), human breast cancer cell line (MDA-MB-231), human colorectal cancer cell line (HCT116), Human ovarian cancer cell line (OVCAR8) | Small Molecule Inhibitor | LC3-II Turnover , p62 Degradation and Rescue | Small Molecule, CQ | Loss of membrane integrity | Cleaved PARP | ApoptosisX | +Autophagy > Cell Death+  -Autophagy > Cell Death- |
| JNK initiates Beclin-1 dependent autophagic cell death against Akt activation | 40 | 10.1016/j.yexcr.2022.113105 | EXPERIMENTAL CELL RESEARCH | Yes | Autophagic Cell Death (ACD) | LC3-II | Cell line, human colorectal cancer cell line (HCT116), "Human colon cancer cell line, Bax/Bak double knockout (HCT116 Bax/Bak KO)", Mouse embryonic fibroblast cell line (MEF) | Small Molecule Inhibitor | LC3-II Turnover | Small Molecule, 3MA | Loss of membrane integrity, Phosphatidyl serine exposure/PI internalization | DNA fragmantation, Phosphatidyl serine exposure | -Apoptosis > Cell Deathx | +Autophagy > Cell Death+  -Autophagy > Cell Death- |
| Elaiophylin Inhibits Tumorigenesis of Human Uveal Melanoma by Suppressing Mitophagy and Inducing Oxidative Stress via Modulating SIRT1/FoxO3a Signaling | 41 | 10.3389/fonc.2022.788496 | FRONTIERS IN ONCOLOGY | Yes | Autophagic Cell Death (ACD) | LC3-II | Cell line, Human uveal melanoma cell line (UM C918), Human uveal melanoma cell line (OCM1A), Human uveal melanoma cell line (Mel270), Human retinal pigment epithelial cell line (ARPE-19) | Small Molecule Inhibitor | No evidence | Not performed | Loss of membrane integrity, Phosphatidyl serine exposure/PI internalization | Phosphatidyl serine exposure | Apoptosis+ | Cell Death+  Autophagy+ |
| High-dose-androgen-induced autophagic cell death to suppress the Enzalutamide-resistant prostate cancer growth via altering the circRNA-BCL2/miRNA-198/AMBRA1 signaling | 42 | 10.1038/s41420-022-00898-6 | CELL DEATH DISCOVERY | Yes | Autophagic Cell Death (ACD) | LC3-II, p62 | Cell line, Human prostate cancer cell line (C4-2 / EnzS-C4-2), Human enzalutamide-resistant prostate cancer cell line (C4-2R / EnzR1-C4-2), Human embryonic kidney cell line (HEK-293T) | Small Molecule Inhibitor | No evidence | Not performed | None | Caspase activation, Cleaved PARP | ApoptosisX | Autophagy+ |
| Photothermal therapy-mediated autophagy in breast cancer treatment: Progress and trends | 43 | 10.1016/j.lfs.2022.120499 | LIFE SCIENCES | No |  |  |  |  |  |  |  |  |  |  |
| Inhibition of Autophagy Facilitates XY03-EA-Mediated Neuroprotection against the Cerebral Ischemia/Reperfusion Injury in Rats | 44 | 10.1155/2022/7013299 | OXIDATIVE MEDICINE AND CELLULAR LONGEVITY | Yes | Autophagic Cell Death (ACD) | LC3-II, p62 | Cell line, PC-12 rat pheochromocytoma cell line, Animal Tissue, Rat brain tissue, Rhesus monkey brain tissue | Small Molecule Inhibitor | No evidence | Not performed | Reduced metabolic activity | None | Unknown | Autophagy+  Viability- |
| Pan-Caspase Inhibitor zVAD Induces Necroptotic and Autophagic Cell Death in TLR3/4-Stimulated Macrophages | 45 | 10.14348/molcells.2021.0193 | MOLECULES AND CELLS | Yes | Autophagic Cell Death (ACD) | LC3-II | Animal Primary Cells, Bone marrow-derived macrophages (BMDMs) | Small Molecule Inhibitor | LC3-II Turnover | Small Molecule, 3MA | Loss of membrane integrity | None | -Apoptosis > Cell Death+ | -Autophagy > Cell Death- |
| Erebosis, a new cell death mechanism during homeostatic turnover of gut enterocytes | 46 | 10.1371/journal.pbio.3001586 | PLOS BIOLOGY | No |  |  |  |  |  |  |  |  |  |  |
| Chemical Characterization of Taif Rose (Rosa damascena Mill var. trigentipetala) Waste Methanolic Extract and Its Hepatoprotective and Antioxidant Effects against Cadmium Chloride (CdCl2)-Induced Hepatotoxicity and Potential Anticancer Activities against Liver Cancer Cells (HepG2) | 47 | 10.3390/cryst12040460 | CRYSTALS | Yes | Autophagic Cell Death (ACD) | Not available | Cell line, human hepatocellular carcinoma (HepG2) | Small Molecule Inhibitor | No evidence | Not performed | Phosphatidyl serine exposure/PI internalization | Phosphatidyl serine exposure | Apoptosis+ | None |
| Necroptosis as a Novel Facet of Mitotic Catastrophe | 48 | 10.3390/ijms23073733 | INTERNATIONAL JOURNAL OF MOLECULAR SCIENCES | No |  |  |  |  |  |  |  |  |  |  |
| Poricoic acid A (PAA) inhibits T-cell acute lymphoblastic leukemia through inducing autophagic cell death and ferroptosis | 49 | 10.1016/j.bbrc.2022.03.105 | BIOCHEMICAL AND BIOPHYSICAL RESEARCH COMMUNICATIONS | Yes | Autophagic Cell Death (ACD) | LC3-II | Cell line, Human T-cell leukemia cell line (Jurkat) | Small Molecule Inhibitor | LC3-II Turnover | Small Molecule, Bafilomycin | None | Caspase activation, Cleaved PARP | -Autophagy > Apoptosis-Apoptosis+ | -Autophagy > Viability+ |
| Regulatory mechanism of alpha-hederin upon cisplatin sensibility in NSCLC at safe dose by destroying GSS/GSH/GPX2 axis-mediated glutathione oxidation-reduction system | 50 | 10.1016/j.biopha.2022.112927 | BIOMEDICINE & PHARMACOTHERAPY | No |  |  |  |  |  |  |  |  |  |  |
| The Dictyostelium Model for Mucolipidosis Type IV | 51 | 10.3389/fcell.2022.741967 | FRONTIERS IN CELL AND DEVELOPMENTAL BIOLOGY | No |  |  |  |  |  |  |  |  |  |  |
| Modulation of energy metabolism to overcome drug resistance in chronic myeloid leukemia cells through induction of autophagy | 52 | 10.1038/s41420-022-00991-w | CELL DEATH DISCOVERY | Yes | Autophagic Cell Death (ACD) | LC3-II | Cell line, Human chronic myeloid leukemia cell line (KBM5), Human imatinib-resistant CML cell line with T315I mutation (KBM5-T315I) | Small Molecule Inhibitor | No evidence | Not performed | Reduced metabolic activity | Phosphatidyl serine exposure | ApoptosisX | Autophagy+  Viability- |
| Anticancer activity of Germacrone terpenoid in human osteosarcoma cells is mediated via autophagy induction, cell cycle disruption, downregulating the cell cycle regulatory protein expressions and cell migration inhibition | 53 | 10.18388/abp.2020_5712 | ACTA BIOCHIMICA POLONICA | Yes | Autophagic Cell Death (ACD) | LC3-II | Cell line, Human osteosarcoma cell line (Saos-2) | Small Molecule Inhibitor | No evidence | Not performed | Reduced metabolic activity | None | Unknown | Autophagy+  Viability- |
| Unraveling the Roles of Protein Kinases in Autophagy: An Updateon Small-Molecule Compounds for Targeted Therapy | 54 | 10.1021/acs.jmedchem.1c02053 | JOURNAL OF MEDICINAL CHEMISTRY | No |  |  |  |  |  |  |  |  |  |  |
| Mogrol suppresses lung cancer cell growth by activating AMPK-dependent autophagic death and inducing p53-dependent cell cycle arrest and apoptosis | 55 | 10.1016/j.taap.2022.116037 | TOXICOLOGY AND APPLIED PHARMACOLOGY | Yes | Autophagic Cell Death (ACD) | LC3-II, p62, Visualization of autophagosomes | Cell line, Human lung adenocarcinoma cell line (A549), Human non-small cell lung cancer cell line (H1299), Human non-small cell lung cancer cell line with EGFR mutation (H1975), Human squamous cell lung carcinoma cell line (SK-MES-1), Human bronchial epithelial cell line (HBE) | Small Molecule Inhibitor | LC3-II Turnover , p62 Degradation and Rescue | Small Molecule, 3MA | Phosphatidyl serine exposure/PI internalization | Caspase activation, Phosphatidyl serine exposure | -Autophagy > Apoptosis-Apoptosis+ | -Autophagy > Viability+ |
| Chemosensitization Effect of Seabuckthorn (Hippophae rhamnoides L.) Pulp Oil via Autophagy and Senescence in NSCLC Cells | 56 | 10.3390/foods11101517 | FOODS | Yes | Autophagic Cell Death (ACD) | LC3-II | Cell line, Human lung adenocarcinoma cell line (A549), Human lung adenocarcinoma cell line (NCI-H23) | Small Molecule Inhibitor | No evidence | Not performed | decreased protein content | Caspase activation | ApoptosisX | Autophagy+  Viability- |
| TNF-like weak inducer of apoptosis / nuclear factor kappa B axis feedback loop promotes spinal cord injury by inducing astrocyte activation | 57 | 10.1080/21655979.2022.2068737 | BIOENGINEERED | Yes | Autophagic Cell Death (ACD) | LC3-II | Animal Tissue, Spinal cord tissue, Animal Primary Cells, Primary mice spinal cord astrocytes and neurons | RNA Interference | No evidence | Not performed | Reduced metabolic activity | None | Unknown | Autophagy+  Viability- |
| LyeTxI-b, a Synthetic Peptide Derived From a Spider Venom, Is Highly Active in Triple-Negative Breast Cancer Cells and Acts Synergistically With Cisplatin | 58 | 10.3389/fmolb.2022.876833 | FRONTIERS IN MOLECULAR BIOSCIENCES | Yes | Autophagic Cell Death (ACD) | Not available | Cell line, human breast cancer cell line (MDA-MB-231), Human embryonic kidney cell line (HEK-293T) | Small Molecule Inhibitor | No evidence | Small Molecule, Bafilomycin | Reduced metabolic activity | None | Unknown | None  Viability- |
| Quercus acutissima Carruth. root extract triggers apoptosis, autophagy and inhibits cell viability in breast cancer cells | 59 | 10.1016/j.jep.2022.115039 | JOURNAL OF ETHNOPHARMACOLOGY | Yes | Autophagic Cell Death (ACD) | LC3-II | Cell line, Human breast cancer cell line (SUM159), human breast cancer cell line (MCF7) | Small Molecule Inhibitor | No evidence | Not performed | None | Caspase activation | Apoptosis+ | Autophagy+ |
| Copper oxide nanoparticles trigger macrophage cell death with misfolding of Cu/Zn superoxide dismutase 1 (SOD1) | 60 | 10.1186/s12989-022-00467-w | PARTICLE AND FIBRE TOXICOLOGY | Yes | Autophagic Cell Death (ACD) | LC3-II, Visualization of autophagosomes | Cell line, Mouse macrophage-like cell line (RAW264.7) | Small Molecule Inhibitor | No evidence | Small Molecule, Bafilomycin | Reduced metabolic activity | None | -Apoptosis > Cell Deathx | -Autophagy > Viability+ |
| Retinal Ganglion Cell Death in Glaucoma: Advances and Caveats | 61 | 10.1080/02713683.2022.2068182 | CURRENT EYE RESEARCH | No |  |  |  |  |  |  |  |  |  |  |
| Modified Titanium Dioxide as a Potential Visible-Light-Activated Photosensitizer for Bladder Cancer Treatment | 62 | 10.1021/acsomega.1c07046 | ACS OMEGA | No |  |  |  |  |  |  |  |  |  |  |
| Melatonin improves arsenic-induced hypertension through the inactivation of the Sirt1/autophagy pathway in rat | 63 | 10.1016/j.biopha.2022.113135 | BIOMEDICINE & PHARMACOTHERAPY | No |  |  |  |  |  |  |  |  |  |  |
| Lithium Enhances Autophagy and Cell Death in Skin Melanoma: An Ultrastructural and Immunohistochemical Study | 64 | 10.1017/S1431927622000745 | MICROSCOPY AND MICROANALYSIS | Yes | Autophagic Cell Death (ACD) | Visualization of autophagosomes | Cell line, mouse skin melanoma cell line (B16), Animal Tissue, Mice skin melanoma tumor tissue | Small Molecule Inhibitor | Autophagolysosome Formation | Not performed | None | Caspase activation, Nuclear margination/ Chromatin condensation, Cell shrinkage | Apoptosis+ | Autophagy+ |
| F-box protein FBXO41 suppresses breast cancer growth by inducing autophagic cell death through facilitating proteasomal degradation of oncogene SKP2 | 65 | 10.1016/j.biocel.2022.106228 | INTERNATIONAL JOURNAL OF BIOCHEMISTRY & CELL BIOLOGY | Yes | Autophagic Cell Death (ACD) | LC3-II, p62 | Cell line, human breast cancer cell line (MCF7), Human embryonic kidney cell line (HEK-293T), human breast cancer cell line (MDA-MB-231), human colorectal cancer cell line (HCT116), "Human colon cancer cell line, p53 knockout (HCT116 p53⁻/⁻)", "Human colon cancer cell line, p21 knockout (HCT116 p21⁻/⁻)", Mouse breast cancer cell line (4T1) | Gene Over-expression | LC3-II Turnover , p62 Degradation and Rescue | Small Molecule, CQ | Reduced metabolic activity, Loss of membrane integrity | Caspase activation | ApoptosisX | -Autophagy > Cell Death-  +Autophagy > Cell Death+ |
| TRIM22 inhibits osteosarcoma progression through destabilizing NRF2 and thus activation of ROS/AMPK/mTOR/autophagy signaling | 66 | 10.1016/j.redox.2022.102344 | REDOX BIOLOGY | Yes | Autophagic Cell Death (ACD), Autophagy Dependent Cell Death | Visualization of autophagosomes, p62 | Cell line, Human osteosarcoma cell line (Saos-2), human osteosarcoma cell line (U-2 OS), Human osteosarcoma cell line (143B), Human osteosarcoma cell line (MG63), Human normal osteoblast cell line (hFOB 1.19), Human embryonic kidney cell line (HEK-293T) | Gene Over-expression | p62 Degradation and Rescue | Small Molecule, 3MA | Phosphatidyl serine exposure/PI internalization | Caspase activation, Phosphatidyl serine exposure | Apoptosis+ | Autophagy+ |
| Transferrin decorated-nanostructured lipid carriers (NLCs) are a promising delivery system for rapamycin in Alzheimer's disease: An in vivo study | 67 | 10.1016/j.bioadv.2022.212827 | BIOMATERIALS ADVANCES | No |  |  |  |  |  |  |  |  |  |  |
| Carnosol Induces p38-Mediated ER Stress Response and Autophagy in Human Breast Cancer Cells | 68 | 10.3389/fonc.2022.911615 | FRONTIERS IN ONCOLOGY | Yes | Autophagic Cell Death (ACD) | LC3-II | Cell line, human breast cancer cell line (MDA-MB-231) | Small Molecule Inhibitor | No evidence | Small Molecule, 3MA, CQ | Loss of membrane integrity | Cleaved PARP | -Apoptosis > Cell Deathx | -Autophagy > Cell Death- |
| Cannabidiol and Cannabigerol Inhibit Cholangiocarcinoma Growth In Vitro via Divergent Cell Death Pathways | 69 | 10.3390/biom12060854 | BIOMOLECULES | Yes | Autophagic Cell Death (ACD) | LC3-II | Cell line, Human intrahepatic cholangiocarcinoma cell line (HuCC-T1), Human extrahepatic cholangiocarcinoma cell line (Mz-ChA-1), Human immortalized non-malignant cholangiocyte cell line (H69) | Small Molecule Inhibitor | No evidence | Not performed | None | Caspase activation | Apoptosis+ | Autophagy+ |
| 6-Gingerol suppresses cell viability, migration and invasion via inhibiting EMT, and inducing autophagy and ferroptosis in LPS-stimulated and LPS-unstimulated prostate cancer cells | 70 | 10.3892/ol.2022.13307 | ONCOLOGY LETTERS | Yes | Autophagic Cell Death (ACD) | LC3-II | Cell line, Human prostate cancer cell line (LNCaP), Human prostate cancer cell line (DU145), Human prostate cancer cell line (PC3) | Small Molecule Inhibitor | No evidence | Small Molecule, PI3K inhibitor | Reduced metabolic activity | None | Unknown | -Autophagy > Viability+ |
| Screening of and mechanism underlying the action of serum- and glucocorticoid-regulated kinase 3-targeted drugs against estrogen receptor-positive breast cancer | 71 | 10.1016/j.ejphar.2022.174982 | EUROPEAN JOURNAL OF PHARMACOLOGY | Yes | Autophagic Cell Death (ACD) | LC3-II, p62 | Cell line, human breast cancer cell line (MCF7), Human breast cancer cell line (ZR-75-1), human breast cancer cell line (T47D), Human non-tumorigenic breast epithelial cell line (MCF-10A) | Small Molecule Inhibitor | No evidence | Small Molecule, 3MA | Reduced metabolic activity | DNA fragmantation | ApoptosisX | -Autophagy > Viability+ |
| Cascade nanozymes based on the butterfly effect for enhanced starvation therapy through the regulation of autophagy | 72 | 10.1039/d2bm00595f | BIOMATERIALS SCIENCE | Yes | Autophagic Cell Death (ACD) | LC3-II, p62, Visualization of autophagosomes | Cell line, Mouse breast cancer cell line (4T1), Human mammary epithelial cell line with SV40 integration (HBL), Human umbilical vein endothelial cell line (HUVEC) | Small Molecule Inhibitor | No evidence | Small Molecule, CQ | Reduced metabolic activity | DNA fragmantation | Apoptosis+ | -Autophagy > Viability+ |
| Death-associated protein kinases and intestinal epithelial homeostasis | 73 | 10.1002/ar.25022 | ANATOMICAL RECORD-ADVANCES IN INTEGRATIVE ANATOMY AND EVOLUTIONARY BIOLOGY | No |  |  |  |  |  |  |  |  |  |  |
| Qili Qiangxin, a compound herbal medicine formula, alleviates hypoxia- reoxygenation-induced apoptotic and autophagic cell death via suppression of ROS/AMPK/mTOR pathway in vitro | 74 | 10.1016/j.joim.2022.04.005 | JOURNAL OF INTEGRATIVE MEDICINE-JIM | Yes | Autophagic Cell Death (ACD) | LC3-II, p62 | Cell line, Rat cardiomyoblast cell line (H9c2) | Physiological stressor | No evidence | Not performed | Phosphatidyl serine exposure/PI internalization, Loss of membrane integrity | Caspase activation, Phosphatidyl serine exposure | Apoptosis+ | Autophagy+  Cell Death+ |
| Autophagy and Programmed Cell Death Are Critical Pathways in Jasmonic Acid Mediated Saline Stress Tolerance in Oryza sativa | 75 | 10.1007/s12010-022-04032-1 | APPLIED BIOCHEMISTRY AND BIOTECHNOLOGY | No |  | Not available |  |  |  | Not performed |  | DNA fragmantation | Unknown | None |
| Pennogenin-3-O-alpha-L-Rhamnopyranosyl-(1 -> 2)-[alpha-L-Rhamnopyranosyl-(1 -> 3)]-beta-D-Glucopyranoside (Spiroconazol A) Isolated from Dioscorea bulbifera L. var. sativa Induces Autophagic Cell Death by p38 MAPK Activation in NSCLC Cells | 76 | 10.3390/ph15070893 | PHARMACEUTICALS | Yes | Autophagic Cell Death (ACD) | LC3-II | Cell line, Human lung adenocarcinoma cell line (A549), Human lung cancer cell line (NCI-H358), human cervical cancer cell line (HeLa), Human cervical cancer cell line (Caski), Human colon cancer cell line (HT-29), human colorectal cancer cell line (HCT116), Human pancreatic cancer cell line (ASPC-1), Human pancreatic cancer cell line (MiaPaCa2), Human normal lung fibroblast cell line (MRC-5) | Small Molecule Inhibitor | LC3-II Turnover | Small Molecule, 3MA | Loss of membrane integrity | Phosphatidyl serine exposure | -Apoptosis > Cell Deathx | -Autophagy > Cell Death- |
| Anticancer Potential of Xanthohumol and Isoxanthohumol Loaded into SBA-15 Mesoporous Silica Particles against B16F10 Melanoma Cells | 77 | 10.3390/ma15145028 | MATERIALS | Yes | Autophagic Cell Death (ACD) | Not available | Cell line, mouse skin melanoma cell line (B16) | Small Molecule Inhibitor | No evidence | Small Molecule, 3MA | Reduced metabolic activity | None | Unknown | None |
| Photothermal effect of albumin-modified gold nanorods diminished neuroblastoma cancer stem cells dynamic growth by modulating autophagy | 78 | 10.1038/s41598-022-15660-2 | SCIENTIFIC REPORTS | Yes | Autophagic Cell Death (ACD) | p62, LC3-II | Cell line, Human neuroblastoma cell line (SH-SY5Y) | Small Molecule Inhibitor | LC3-II Turnover , p62 Degradation and Rescue | Small Molecule, CQ | Reduced metabolic activity, Loss of membrane integrity | Phosphatidyl serine exposure | Apoptosis+ | Autophagy+ |
| Lomitapide, a cholesterol-lowering drug, is an anticancer agent that induces autophagic cell death via inhibiting mTOR | 79 | 10.1038/s41419-022-05039-6 | CELL DEATH & DISEASE | Yes | Autophagic Cell Death (ACD) | LC3-II | Cell line, "Human colon cancer cell line (HCT116, p53 wild-type)", "Human colon cancer cell line (HT29, p53 mutant)", "Human colon cancer cell line (SW480, p53 mutant)", human breast cancer cell line (MDA-MB-231), human breast cancer cell line (MDA-MB-468), Human melanoma cell line (A375), Human melanoma cell line (A2058), Human normal colon mucosa cell line (NCM460), Human stomach cancer cell line (HS746T), Human stomach cancer cell line (SNU1), Human stomach cancer cell line (SNU216) | Small Molecule Inhibitor | No evidence | Both, 3MA | Reduced metabolic activity | Caspase activation | ApoptosisX | -Autophagy > Viability+ |
| KS10076, a chelator for redox-active metal ions, induces ROS-mediated STAT3 degradation in autophagic cell death and eliminates ALDH1(+) stem cells | 80 | 10.1016/j.celrep.2022.111077 | CELL REPORTS | Yes | Autophagic Cell Death (ACD) | LC3-II | Human Primary Cells, Animal Tissue, Mouse xenograft tumor tissue, Cell line, human colorectal cancer cell line (HCT116), human colon adenocarcinoma cell line (SW480), Human colon cancer cell line (COLO205), Patient-derived colorectal cancer organoids (PDOs) | Small Molecule Inhibitor | No evidence | Genetic | Reduced metabolic activity | Bcl-2 expression | -Apoptosis > ViabilityX | -Autophagy > Viability+ |
| An Endogenous H2S-Activated Nanoplatform for Triple Synergistic Therapy of Colorectal Cancer | 81 | 10.1021/acs.nanolett.2c01346 | NANO LETTERS | Yes | Autophagic Cell Death (ACD) | LC3-II, p62 | Cell line, human colorectal cancer cell line (HCT116) | Small Molecule Inhibitor | No evidence | Not performed | Reduced metabolic activity | Caspase activation | Apoptosis+ | Autophagy+  Cell Death+ |
| Berberine Induces Autophagic Cell Death by Inactivating the Akt/mTOR Signaling Pathway | 82 | 10.1055/a-1752-0311 | PLANTA MEDICA | Yes | Autophagic Cell Death (ACD) | LC3-II, Visualization of autophagosomes | Cell line, mouse skin melanoma cell line (B16) | Small Molecule Inhibitor | LC3-II Turnover | Small Molecule, 3MA | Reduced metabolic activity | BAX expression | Unknown | -Autophagy > Viability+ |
| Diterpene promptly executes a non-canonical autophagic cell death in doxorubicin-resistant lung cancer | 83 | 10.1016/j.biopha.2022.113443 | BIOMEDICINE & PHARMACOTHERAPY | Yes | Autophagic Cell Death (ACD) | LC3-II, p62 | Cell line, "Human lung cancer cell line (A549, Dox-S)", "Human lung cancer cell line (A549, Dox-R, doxorubicin-resistant derivative)" | Small Molecule Inhibitor | No evidence | Small Molecule, Bafilomycin | Reduced metabolic activity | DNA fragmantation, Cleaved PARP | Apoptosis+ | -Autophagy > Viability+ |
| The effects of glutamine supplementation on markers of apoptosis and autophagy in sickle cell disease peripheral blood mononuclear cells | 84 | 10.1016/j.ctim.2022.102856 | COMPLEMENTARY THERAPIES IN MEDICINE | Yes | Autophagic Cell Death (ACD) | LC3-II | Human Primary Cells, Peripheral blood mononuclear cells (PBMCs) | Physiological stressor, Small Molecule Inhibitor | No evidence | Not performed | None | BAX expression | Unknown | Autophagy+ |
| Polyphyllin I Promotes Autophagic Cell Death and Apoptosis of Colon Cancer Cells via the ROS-Inhibited AKT/mTOR Pathway | 85 | 10.3390/ijms23169368 | INTERNATIONAL JOURNAL OF MOLECULAR SCIENCES | Yes | Autophagic Cell Death (ACD) | LC3-II | Cell line, human colon adenocarcinoma cell line (SW480) | Small Molecule Inhibitor | LC3-II Turnover | Small Molecule, CQ | Reduced metabolic activity | Nuclear margination/ Chromatin condensation | -Autophagy > Apoptosis-Apoptosis+ | -Autophagy > Viability+ |
| Diindolylmethane Inhibits Cadmium-Induced Autophagic Cell Death via Regulation of Oxidative Stress in HEL299 Human Lung Fibroblasts | 86 | 10.3390/molecules27165215 | MOLECULES | Yes | Autophagic Cell Death (ACD) | p62, LC3-II | Cell line, Human embryonic lung cell line (HEL 299) | Small Molecule Inhibitor | No evidence | Small Molecule, Bafilomycin | Reduced metabolic activity | Phosphatidyl serine exposure | ApoptosisX | -Autophagy > Cell Death- |
| Adenosine derivatives from Cordyceps exert antitumor effects against ovarian cancer cells through ENT1-mediated transport, induction of AMPK signaling, and consequent autophagic cell death | 87 | 10.1016/j.biopha.2022.113491 | BIOMEDICINE & PHARMACOTHERAPY | Yes | Autophagic Cell Death (ACD) | LC3-II, Visualization of autophagosomes | Cell line, Human ovarian cancer cell line (A2780), Human ovarian cancer cell line (OVCAR3), Human ovarian cancer cell line (SKOV3), Human ovarian cancer cell line (TOV112D), Human embryonic kidney cell line (HEK293) | Small Molecule Inhibitor | Autophagolysosome Formation | Not performed | Reduced metabolic activity | Phosphatidyl serine exposure | ApoptosisX | Autophagy+  Viability- |
| Carbon dots supported single Fe atom nanozyme for drug-resistant glioblastoma therapy by activating autophagy-lysosome pathway | 88 | 10.1016/j.nantod.2022.101530 | NANO TODAY | Yes | Autophagic Cell Death (ACD) | LC3-II, p62 | Cell line, Human glioblastoma cell line (U87MG), "Human glioblastoma cell line (U251-TR, temozolomide-resistant)" | Small Molecule Inhibitor | No evidence | Not performed | Loss of membrane integrity, Phosphatidyl serine exposure/PI internalization, Reduced metabolic activity | DNA fragmantation, Caspase activation, Phosphatidyl serine exposure | Apoptosis+ | Autophagy+  Cell Death+ |
| Dysregulated sphingolipid metabolism and autophagy in granulosa cells of women with endometriosis | 89 | 10.3389/fendo.2022.906570 | FRONTIERS IN ENDOCRINOLOGY | Yes | Autophagic Cell Death (ACD) | Not available | Human Primary Cells, Human cumulus cells (CCs), Human mural granulosa cells (MGCs) | Small Molecule Inhibitor | No evidence | Not performed | None | None | Unknown | None |
| FOXA1 prevents nutrients deprivation induced autophagic cell death through inducing loss of imprinting of IGF2 in lung adenocarcinoma | 90 | 10.1038/s41419-022-05150-8 | CELL DEATH & DISEASE | Yes | Autophagic Cell Death (ACD) | LC3-II, Visualization of autophagosomes | Cell line, Human lung adenocarcinoma cell line (A549), Human lung adenocarcinoma cell line (Calu-3), Human lung adenocarcinoma cell line (PC-9) | Gene Over-expression | LC3-II Turnover | Small Molecule, 3MA, CQ | Reduced colony formation | None | -Apoptosis > ViabilityX | -Autophagy > Viability+ |
| Novel cardioprotective mechanism for Empagliflozin in nondiabetic myocardial infarction with acute hyperglycemia | 91 | 10.1016/j.biopha.2022.113606 | BIOMEDICINE & PHARMACOTHERAPY | Yes | Autosis, Autophagic Cell Death (ACD) | LC3-II, p62 | Animal Primary Cells, Human Primary Cells, Animal Tissue, Neonatal rat ventricular myocytes (NRVMs), Heart tissue from wild-type C57BL/6J mice, Human induced pluripotent stem cell-derived cardiomyocytes (hiPSC-CMs) | Small Molecule Inhibitor | No evidence | Not performed | None | None | Unknown | Autophagy+ |
| Iron metabolism mediates microglia susceptibility in ferroptosis | 92 | 10.3389/fncel.2022.995084 | FRONTIERS IN CELLULAR NEUROSCIENCE | Yes | Autophagic Cell Death (ACD) | Not available | Animal Primary Cells, Primary rat cortical astrocytes, Primary rat cortical microglia, Primary rat cortical neurons | Small Molecule Inhibitor | No evidence | Not performed | Reduced metabolic activity, Loss of membrane integrity | None | Unknown | None, Viability- |
| Overexpression of TRIM32 promotes pancreatic beta-cell autophagic cell death through Akt/mTOR pathway under high glucose conditions | 93 | 10.1002/cbin.11897 | CELL BIOLOGY INTERNATIONAL | Yes | Autophagic Cell Death (ACD) | LC3-II | Cell line, Rat insulinoma cell line (INS-1) | Gene Over-expression | No evidence | Not performed | Loss of membrane integrity | DNA fragmantation | Apoptosis+ | Autophagy+  Cell Death+ |
| Quercetin induces autophagy-associated death in HL-60 cells through CaMKK beta/AMPK/mTOR signal pathway | 94 | 10.3724/abbs.2022117 | ACTA BIOCHIMICA ET BIOPHYSICA SINICA | Yes | Autophagic Cell Death (ACD), Autophagy Associated Cell Death (AACD) | LC3-II, p62, Visualization of autophagosomes | Cell line, Human promyelocytic leukemia cell line (HL-60), Human acute myeloid leukemia cell line (KG-1), Human acute myeloid leukemia cell line (THP-1) | Small Molecule Inhibitor | LC3-II Turnover , p62 Degradation and Rescue | Small Molecule, 3MA | Reduced colony formation | Phosphatidyl serine exposure | Apoptosis+ | -Autophagy > Cell Death- |
| Overcoming Basal Autophagy, Kangai Injection Enhances Cisplatin Cytotoxicity by Regulating FOXO3a-Dependent Autophagic Cell Death and Apoptosis in Human Lung Adenocarcinoma A549/DDP Cells | 95 | 10.1155/2022/6022981 | BIOMED RESEARCH INTERNATIONAL | Yes | Autophagic Cell Death (ACD) | LC3-II, p62 | Cell line, Human lung adenocarcinoma cell line (A549), Human cisplatin-resistant lung adenocarcinoma cell line (A549/DDP) | Gene Over-expression | No evidence | Small Molecule, Bafilomycin | Reduced metabolic activity | Caspase activation, BAX expression, Phosphatidyl serine exposure | -Apoptosis > Viability+ | -Autophagy > Viability+ |
| The Yin and Yang dualistic features of autophagy in thermal burn wound healing | 96 | 10.1177/03946320221125090 | INTERNATIONAL JOURNAL OF IMMUNOPATHOLOGY AND PHARMACOLOGY | No |  |  |  |  |  |  |  |  |  |  |
| Quantification of ferroptosis pathway status revealed heterogeneity in breast cancer patients with distinct immune microenvironment | 97 | 10.3389/fonc.2022.956999 | FRONTIERS IN ONCOLOGY | No |  |  |  |  |  |  |  |  |  |  |
| Targeting glucose-6-phosphate dehydrogenase by 6-AN induces ROS-mediated autophagic cell death in breast cancer | 98 | 10.1111/febs.16614 | FEBS JOURNAL | Yes | Autophagic Cell Death (ACD) | LC3-II, p62 | Cell line, human breast cancer cell line (MDA-MB-231), human breast cancer cell line (MCF7), Human breast cancer cell line (SKBR3), Human embryonic kidney cell line (HEK-293T), Human non-tumorigenic breast epithelial cell line (MCF-10A) | Small Molecule Inhibitor | LC3-II Turnover , p62 Degradation and Rescue | Small Molecule, CQ | Reduced metabolic activity, Loss of membrane integrity | Caspase activation, Cleaved PARP, Phosphatidyl serine exposure | ApoptosisX | -Autophagy > Cell Death- |
| Engineering mitochondrial uncoupler synergistic photodynamic nanoplatform to harness immunostimulatory pro-death autophagy/ mitophagy | 99 | 10.1016/j.biomaterials.2022.121796 | BIOMATERIALS | Yes | Autophagic Cell Death (ACD) | LC3-II, p62 | Cell line, Mouse breast cancer cell line (4T1) | Small Molecule Inhibitor | No evidence | Small Molecule, 3MA, CQ | Reduced metabolic activity | Phosphatidyl serine exposure | Apoptosis+ | -Autophagy > Cell Death- |
| Cellular Hypoxia Mitigation by Dandelion-like Nanoparticles for Synergistic Photodynamic Therapy of Oral Squamous Cell Carcinoma | 100 | 10.1021/acsami.2c10021 | ACS APPLIED MATERIALS & INTERFACES | No |  |  |  |  |  |  |  |  |  |  |
| The natural medicinal fungus Huaier promotes the anti-hepatoma efficacy of sorafenib through the mammalian target of rapamycin-mediated autophagic cell death | 101 | 10.1007/s12032-022-01797-7 | MEDICAL ONCOLOGY | Yes | Autophagic Cell Death (ACD) | LC3-II, p62, Visualization of autophagosomes | Cell line, Human hepatocellular carcinoma cell line (Hep3B), Human hepatocellular carcinoma cell line (Huh7) | Small Molecule Inhibitor | Autophagolysosome Formation | Not performed | Reduced metabolic activity, Reduced colony formation | Phosphatidyl serine exposure | Apoptosis+ | Autophagy+  Viability- |
| Diclofenac Sensitizes Signet Ring Cell Gastric Carcinoma Cells to Cisplatin by Activating Autophagy and Inhibition of Survival Signal Pathways | 102 | 10.3390/ijms232012066 | INTERNATIONAL JOURNAL OF MOLECULAR SCIENCES | Yes | Autophagic Cell Death (ACD) | LC3-II, Visualization of autophagosomes | Cell line, Human gastric cancer cisplatin-sensitive cell line (KATO III), Human gastric cancer cisplatin-resistant cell line (KATO/DDP) | Small Molecule Inhibitor | No evidence | Small Molecule, 3MA | Loss of membrane integrity | Phosphatidyl serine exposure, Cleaved PARP | ApoptosisX | -Autophagy > Cell Death- |
| Selaginella tamariscina Inhibits Glutamate-Induced Autophagic Cell Death by Activating the PI3K/AKT/mTOR Signaling Pathways | 103 | 10.3390/ijms231911445 | INTERNATIONAL JOURNAL OF MOLECULAR SCIENCES | Yes | Autophagic Cell Death (ACD) | LC3-II, p62 | Cell line, Mouse hippocampal neuronal cell line (HT22) | Small Molecule Inhibitor | LC3-II Turnover , p62 Degradation and Rescue | Small Molecule, CQ | Loss of membrane integrity, Reduced metabolic activity | Phosphatidyl serine exposure, Cleaved PARP | Apoptosis+ | Autophagy+  Cell Death+ |
| 6,6 '-((Methylazanedyl)bis(methylene))bis(2,4-dimethylphenol) Induces Autophagic Associated Cell Death through mTOR-Mediated Autophagy in Lung Cancer | 104 | 10.3390/molecules27196230 | MOLECULES | Yes | Autophagy Associated Cell Death (AACD), Autophagic Cell Death (ACD) | LC3-II, p62, Visualization of autophagosomes | Cell line, Human lung adenocarcinoma cell line (A549) | Small Molecule Inhibitor | Autophagolysosome Formation | Not performed | Reduced metabolic activity, Phosphatidyl serine exposure/PI internalization | Caspase activation, DNA fragmantation, Phosphatidyl serine exposure | Apoptosis+ | +Autophagy > Cell Death+ |
| Enhanced Cytotoxic Effects of Arenite in Combination with Active Bufadienolide Compounds against Human Glioblastoma Cell Line U-87 | 105 | 10.3390/molecules27196577 | MOLECULES | Yes | Autophagic Cell Death (ACD) | LC3-II | Cell line, Human glioblastoma cell line (U87MG) | Small Molecule Inhibitor | No evidence | Not performed | Reduced metabolic activity, Loss of membrane integrity | Phosphatidyl serine exposure, Caspase activation | Apoptosis+ | Autophagy+  Cell Death+ |
| Diprenylated flavonoids from licorice induce death of SW480 colorectal cancer cells by promoting autophagy: Activities of lupalbigenin and 6,8-diprenylgenistein | 106 | 10.1016/j.jep.2022.115488 | JOURNAL OF ETHNOPHARMACOLOGY | Yes | Autophagic Cell Death (ACD) | LC3-II, p62 | Cell line, "Human colon cancer cell line (SW480, p53 mutant)" | Small Molecule Inhibitor | LC3-II Turnover , p62 Degradation and Rescue | Small Molecule, CQ | Reduced metabolic activity | Phosphatidyl serine exposure, Caspase activation | -Autophagy > Apoptosis-Apoptosis+ | -Autophagy > Viability+ |
| Biologically synthesized black ginger-selenium nanoparticle induces apoptosis and autophagy of AGS gastric cancer cells by suppressing the PI3K/Akt/mTOR signaling pathway | 107 | 10.1186/s12951-022-01576-6 | JOURNAL OF NANOBIOTECHNOLOGY | Yes | Autophagic Cell Death (ACD) | LC3-II, p62, Visualization of autophagosomes | Cell line, Human gastric cancer cell line (AGS), Human keratinocyte cell line (HaCaT) | Small Molecule Inhibitor | LC3-II Turnover , p62 Degradation and Rescue | Small Molecule, CQ | Reduced metabolic activity | Phosphatidyl serine exposure, Caspase activation, DNA fragmantation | Apoptosis+ | Viability-  Autophagy+ |
| Emodin coupled with high LET neutron beam-a novel approach to treat on glioblastoma | 108 | 10.1093/jrr/rrac061 | JOURNAL OF RADIATION RESEARCH | Yes | Autophagic Cell Death (ACD) | Visualization of autophagosomes | Cell line, Human glioma cell line (LN18), Human glioma cell line (LN428) | Small Molecule Inhibitor | No evidence | Not performed | Reduced metabolic activity, Phosphatidyl serine exposure/PI internalization | Cleaved PARP, Caspase activation, Phosphatidyl serine exposure, DNA fragmantation | Apoptosis+ | Autophagy+  Viability- |
| Chinese dragon's blood ethyl acetate extract suppresses gastric cancer progression through induction of apoptosis and autophagy mediated by activation of MAPK and downregulation of the mTOR-Beclin1 signalling cascade | 109 | 10.1002/ptr.7652 | PHYTOTHERAPY RESEARCH | Yes | Autophagic Cell Death (ACD) | LC3-II, p62 | Cell line, Human gastric cancer cell line (MGC-803), Human gastric adenocarcinoma cell line (HGC27), Human normal gastric epithelial cell line (GES-1) | Small Molecule Inhibitor | No evidence | Small Molecule, CQ | Reduced metabolic activity, Reduced colony formation, Phosphatidyl serine exposure/PI internalization | Cleaved PARP, Phosphatidyl serine exposure | -Apoptosis > Viability+ | -Autophagy > Viability+ |
| Larotrectinib induces autophagic cell death through AMPK/mTOR signalling in colon cancer | 110 | 10.1111/jcmm.17530 | JOURNAL OF CELLULAR AND MOLECULAR MEDICINE | Yes | Autophagy Associated Cell Death (AACD), Autophagic Cell Death (ACD) | LC3-II, p62, Visualization of autophagosomes | Cell line, Human colon cancer cell line (COLO205), "Human colon cancer cell line (HCT116, p53 wild-type)" | Small Molecule Inhibitor | LC3-II Turnover , p62 Degradation and Rescue | Both, CQ | Reduced metabolic activity | None | Unknown | Autophagy+  Viability- |
| Hernandezine induces autophagic cell death in human pancreatic cancer cells via activation of the ROS/AMPK signaling pathway | 111 | 10.1038/s41401-022-01006-1 | ACTA PHARMACOLOGICA SINICA | Yes | Autophagic Cell Death (ACD) | LC3-II, p62 | Cell line, Human pancreatic cancer cell line (Capan-1), Human pancreatic cancer cell line (SW1990) | Small Molecule Inhibitor | LC3-II Turnover , p62 Degradation and Rescue | Both, Bafilomycin, CQ | Reduced metabolic activity, Phosphatidyl serine exposure/PI internalization | Phosphatidyl serine exposure | -Autophagy > Apoptosis-Apoptosis+ | Cell Death+  Autophagy+ |
| Branched-chain amino acids (BCAA) administration increases autophagy and the autophagic pathway in brain tissue of rats submitted to a Maple Syrup Urine Disease (MSUD) protocol | 112 | 10.1007/s11011-022-01109-y | METABOLIC BRAIN DISEASE | Yes | Autophagic Cell Death (ACD) | LC3-II | Animal Tissue, Rat brain tissue | Small Molecule Inhibitor | No evidence | Not performed | None | None | Unknown | Autophagy+ |
| A novel BH3 mimetic Bcl-2 inhibitor promotes autophagic cell death and reduces in vivo Glioblastoma tumor growth | 113 | 10.1038/s41420-022-01225-9 | CELL DEATH DISCOVERY | Yes | Autophagic Cell Death (ACD) | LC3-II, Visualization of autophagosomes | Cell line, Human glioblastoma cell line (U87MG), Human glioblastoma cell line (A172), Human glioma cell line (LN18), Human glioblastoma cell line (YKG1), Human neuroblastoma cell line (SH-SY5Y), Human umbilical vein endothelial cell line (HUVEC), Human promyelocytic leukemia cell line (HL-60) | Small Molecule Inhibitor | No evidence | Not performed | Reduced colony formation | Cleaved PARP, Caspase activation, Phosphatidyl serine exposure | ApoptosisX | Autophagy+  Viability- |
| Phytochemical Profile of the Ethanol Extract of Malvaviscus arboreus Red Flower and Investigation of the Antioxidant, Antimicrobial, and Cytotoxic Activities | 114 | 10.3390/antibiotics11111652 | ANTIBIOTICS-BASEL | Yes | Autophagic Cell Death (ACD) | Visualization of autophagosomes | Cell line, human hepatocellular carcinoma (HepG2) | Small Molecule Inhibitor | No evidence | Not performed | Phosphatidyl serine exposure/PI internalization, decreased protein content | Phosphatidyl serine exposure, Caspase activation, DNA fragmantation | Apoptosis+ | Autophagy+  Cell Death+ |
| Mallotucin D, a Clerodane Diterpenoid from Croton crassifolius, Suppresses HepG2 Cell Growth via Inducing Autophagic Cell Death and Pyroptosis | 115 | 10.3390/ijms232214217 | INTERNATIONAL JOURNAL OF MOLECULAR SCIENCES | Yes | Autophagic Cell Death (ACD) | LC3-II, p62, Visualization of autophagosomes | Cell line, human hepatocellular carcinoma (HepG2) | Small Molecule Inhibitor | Autophagolysosome Formation | Not performed | Reduced metabolic activity, Loss of membrane integrity | Caspase activation, Phosphatidyl serine exposure | Apoptosis+ | Autophagy+  Cell Death+ |
| Colon cancer therapy with calcium phosphate nanoparticles loading bioactive compounds from Euphorbia lathyris: In vitro and in vivo assay | 116 | 10.1016/j.biopha.2022.113723 | BIOMEDICINE & PHARMACOTHERAPY | Yes | Autophagic Cell Death (ACD) | Visualization of autophagosomes | Cell line, Human colon cancer cell line (T84), Mouse colon cancer cell line (MC-38), human normal colon cell line (CCD-18Co) | Small Molecule Inhibitor | Autophagolysosome Formation | Not performed | None | Nuclear margination/ Chromatin condensation | Apoptosis+ | Autophagy+ |
| Ciclopirox drives growth arrest and autophagic cell death through STAT3 in gastric cancer cells | 117 | 10.1038/s41419-022-05456-7 | CELL DEATH & DISEASE | Yes | Autophagic Cell Death (ACD) | LC3-II, p62 | Cell line, Human gastric cancer cell line (AGS), Human gastric cancer cell line (SGC-7901), Human gastric cancer cell line (MGC-803), Human normal gastric epithelial cell line (GES-1) | Small Molecule Inhibitor | LC3-II Turnover , p62 Degradation and Rescue | Small Molecule, 3MA, CQ, Bafilomycin | Reduced colony formation, Reduced metabolic activity, Loss of membrane integrity | Phosphatidyl serine exposure | ApoptosisX | -Autophagy > Cell Death-  +Autophagy > Cell Death+ |
| CDK4/6 Inhibition Enhances the Efficacy of Standard Chemotherapy Treatment in Malignant Pleural Mesothelioma Cells | 118 | 10.3390/cancers14235925 | CANCERS | Yes | Autophagic Cell Death (ACD) | LC3-II, Visualization of autophagosomes | Cell line, Human malignant pleural mesothelioma cell line (MSTO-211H), Human malignant pleural mesothelioma cell line (H28) | Small Molecule Inhibitor | No evidence | Small Molecule, 3MA | Reduced colony formation, Reduced metabolic activity, Phosphatidyl serine exposure/PI internalization | Cleaved PARP, Caspase activation, Phosphatidyl serine exposure | -Autophagy > Apoptosis-Apoptosis+ | Cell Death+  Autophagy+ |
| Beyond Traditional Use of Alchemilla vulgaris: Genoprotective and Antitumor Activity In Vitro | 119 | 10.3390/molecules27238113 | MOLECULES | Yes | Autophagic Cell Death (ACD) | Not available | Cell line, human breast cancer cell line (MCF7), Human melanoma cell line (A375), Human lung adenocarcinoma cell line (A549), human colorectal cancer cell line (HCT116) | Small Molecule Inhibitor | No evidence | Small Molecule, 3MA | decreased protein content, Reduced metabolic activity, Loss of membrane integrity | Phosphatidyl serine exposure | Unknown | None |
| Resveratrol Inhibits Proliferation and Induces Autophagy by Blocking SREBP1 Expression in Oral Cancer Cells | 120 | 10.3390/molecules27238250 | MOLECULES | Yes | Autophagic Cell Death (ACD) | LC3-II, p62 | Cell line, "Human oral squamous cell carcinoma cell line (HSC-2, 3, 4)", Human oral squamous cell carcinoma cell line (Ca9-22), Human oral squamous cell carcinoma cell line (SAS) | Small Molecule Inhibitor | No evidence | Small Molecule, 3MA | Reduced metabolic activity | Caspase activation | Apoptosis+ | -Autophagy > Viability+ |
| Polystyrene microplastic particles induce autophagic cell death in BEAS-2B human bronchial epithelial cells | 121 | 10.1002/tox.23705 | ENVIRONMENTAL TOXICOLOGY | Yes | Autophagic Cell Death (ACD) | p62, Visualization of autophagosomes | Cell line, Human bronchial epithelial cell line (BEAS-2B) | Small Molecule Inhibitor | Autophagolysosome Formation | Genetic | None | Nuclear margination/ Chromatin condensation | Apoptosis+ | Autophagy+ |
| Role of ATP5G3 in sodium nitroprusside-induced cell death in cervical carcinoma cells | 122 | 10.1002/jbt.23267 | JOURNAL OF BIOCHEMICAL AND MOLECULAR TOXICOLOGY | Yes | Autophagic Cell Death (ACD) | p62 | Cell line, human cervical cancer cell line (HeLa) | Small Molecule Inhibitor | No evidence | Bafilomycin, Small Molecule, Concanamycin-A | Reduced metabolic activity | None | -Apoptosis > ViabilityX | -Autophagy > Viability+ |
| Gracillin exerts anti-melanoma effects in vitro and in vivo: role of DNA damage, apoptosis and autophagy | 123 | 10.1016/j.phymed.2022.154526 | PHYTOMEDICINE | Yes | Autophagic Cell Death (ACD) | LC3-II | Cell line, mouse skin melanoma cell line (B16), Human melanoma cell line (A375) | Small Molecule Inhibitor | LC3-II Turnover | Small Molecule, 3MA, CQ, Bafilomycin | Reduced metabolic activity, Reduced colony formation, Phosphatidyl serine exposure/PI internalization | Cleaved PARP, Phosphatidyl serine exposure | Apoptosis+ | +Autophagy > Cell Death+  -Autophagy > Cell Death- |
| L-Carnitine Mitigates Trazadone Induced Rat Cardiotoxicity Mediated via Modulation of Autophagy and Oxidative Stress | 124 | 10.1007/s12012-022-09759-1 | ACTA BIOCHIMICA ET BIOPHYSICA SINICA | No |  | LC3-II |  |  |  | Not performed | None | None | Unknown | None |
| Centrifugally concentric ring-patterned drug-loaded polymeric coating as an intraocular lens surface modification for efficient prevention of posterior capsular opacification | 125 | 10.1016/j.actbio.2021.11.018 | ONCOLOGY LETTERS | Yes | Autophagic Cell Death (ACD), Autophagy Mediated Cell Death | LC3-II | Cell line, HLEC-B3 human lens epithelial cell line, Animal Tissue, Rabbit eye tissue | Small Molecule Inhibitor | No evidence | Not performed | Loss of membrane integrity, Reduced metabolic activity, Phosphatidyl serine exposure/PI internalization | Phosphatidyl serine exposure | Apoptosis+ | Autophagy+  Cell Death+ |
| Ablation of FUNDC1-dependent mitophagy renders myocardium resistant to paraquat-induced ferroptosis and contractile dysfunction | 126 | 10.1016/j.bbadis.2022.166448 | BIOMOLECULES & THERAPEUTICS | No |  |  |  |  |  |  |  |  |  |  |
| Induction of tumor cell autosis by myxoma virus-infected CAR-T and TCR-T cells to overcome primary and acquired resistance | 127 | 10.1016/j.ccell.2022.08.001 | ASIAN PACIFIC JOURNAL OF TROPICAL BIOMEDICINE | No |  |  |  |  |  |  |  |  |  |  |
| Leveraging diverse cell-death patterns to predict the prognosis and drug sensitivity of triple-negative breast cancer patients after surgery | 128 | 10.1016/j.ijsu.2022.106936 | ARCHIVES OF MEDICAL SCIENCE | No |  |  |  |  |  |  |  |  |  |  |
| Geraniin inhibits cell growth and promoted autophagy-mediated cell death in the nasopharyngeal cancer C666-1 cells | 129 | 10.1016/j.sjbs.2021.08.076 | INTERNATIONAL JOURNAL OF MEDICAL SCIENCES | Yes | Autophagy Mediated Cell Death | LC3-II | Cell line, Human nasopharyngeal carcinoma cell line (C666-1) | Small Molecule Inhibitor | No evidence | Not performed | Reduced metabolic activity, Loss of membrane integrity | None | Unknown | Autophagy+  Cell Death+ |
| Recovery Mechanism of Endoplasmic Reticulum Revealed by Fluorescence Lifetime Imaging in Live Cells | 130 | 10.1021/acs.analchem.2c00216 | JOURNAL OF CANCER | No |  |  |  |  |  |  |  |  |  |  |
| Targeting Regulated Cell Death with Pharmacological Small Molecules: An Update on Autophagy-Dependent Cell Death, Ferroptosis, and Necroptosis in Cancer | 131 | 10.1021/acs.jmedchem.1c01572 | MOLECULES AND CELLS | No |  |  |  |  |  |  |  |  |  |  |
| XDeathDB: a visualization platform for cell death molecular interactions | 132 | 10.1038/s41419-021-04397-x | QUALITY ASSURANCE AND SAFETY OF CROPS & FOODS | No |  |  |  |  |  |  |  |  |  |  |
| PGK1 represses autophagy-mediated cell death to promote the proliferation of liver cancer cells by phosphorylating PRAS40 | 133 | 10.1038/s41419-022-04499-0 | EXCLI JOURNAL | Yes | Autophagy Mediated Cell Death | p62, LC3-II, Visualization of autophagosomes | Cell line, Human Ewing’s sarcoma cell line (A673), human hepatocellular carcinoma (HepG2), Human liver cancer cell line (SNU449), Human embryonic kidney cell line (HEK-293T) | Gene Over-expression | No evidence | Small Molecule, 3MA, CQ, Non specific Small Molecule, PRAS40 | Phosphatidyl serine exposure/PI internalization, Reduced colony formation | Phosphatidyl serine exposure, Cleaved PARP | -Apoptosis > Cell Death- | -Autophagy > Cell Death- |
| Induction of apoptosis and autosis in cardiomyocytes by the combination of homocysteine and copper via NOX-mediated p62 expression | 134 | 10.1038/s41420-022-00870-4 | ACTA BIOCHIMICA POLONICA | Yes | Autophagic Cell Death (ACD), Autosis | LC3-II, p62 | Cell line, Rat cardiomyoblast cell line (H9c2), Animal Primary Cells, Neonatal rat ventricular myocytes (NRVMs), Animal Tissue, Rat heart tissue | Small Molecule Inhibitor | LC3-II Turnover , p62 Degradation and Rescue | Both, CQ | Loss of membrane integrity, Phosphatidyl serine exposure/PI internalization | Caspase activation, Phosphatidyl serine exposure | -Autophagy > Apoptosis  -Apoptosis > Autophagy+  -Apoptosis > Cell Deathx | -Autophagy > Cell Death- |
| Extracellular SQSTM1 exacerbates acute pancreatitis by activating autophagy-dependent ferroptosis | 135 | 10.1080/15548627.2022.2152209 | ANTICANCER RESEARCH | No |  |  |  |  |  |  |  |  |  |  |
| STAT3 Enhances Sensitivity of Glioblastoma to Drug-Induced Autophagy-Dependent Cell Death | 136 | 10.3390/cancers14020339 | CCS CHEMISTRY | Yes | Autophagy Dependent Cell Death | LC3-II, Visualization of autophagosomes | Cell line, Human glioblastoma cell line (A172), "Human glioblastoma cell line (MZ-18, MZ-54, MZ-256, MZ-304)", Human glioblastoma cell line (U87MG), Human glioblastoma cell line (U251-MG), Human glioblastoma cell line (U343-MG), Human glioblastoma cell line (U373-MG), Human glioblastoma cell line (LN229), Murine glioblastoma cell line (Tu2449), Murine glioblastoma cell line (Tu-9648), Human embryonic kidney cell line (HEK-293T), "Glioblastoma Stem Cell Line (PB1, FPW1, MN1, RKI1, RN1, MMK1)", "Glioblastoma Stem Cell Line (NCH1425, NCH601, NCH644, NCH421k)" | Small Molecule Inhibitor | Autophagolysosome Formation | Not performed | Reduced metabolic activity, Phosphatidyl serine exposure/PI internalization | Phosphatidyl serine exposure | Apoptosis+ | Autophagy+  Cell Death+ |
| Anticolon Cancer Effect of Korean Red Ginseng via Autophagy- and Apoptosis-Mediated Cell Death | 137 | 10.3390/nu14173558 | ARCHIVES OF PHARMACY PRACTICE | Yes | Autophagy Mediated Cell Death | LC3-II, Visualization of autophagosomes | Cell line, human colorectal cancer cell line (HCT116), Human colon cancer cell line (SNU-1033), Human colon cancer cell line (Caco-2), "Human colon cancer cell line (HT29, p53 mutant)", Normal human colon cell line (FHC) | Small Molecule Inhibitor | LC3-II Turnover | Both, Bafilomycin | Phosphatidyl serine exposure/PI internalization, Reduced metabolic activity | Phosphatidyl serine exposure, Caspase activation, Cleaved PARP, DNA fragmantation | -Apoptosis > Cell Death- | -Autophagy > Viability+ |
